# KaryoCreate: a new CRISPR-based technology to generate chromosome-specific aneuploidy by targeting human centromeres

**DOI:** 10.1101/2022.09.27.509580

**Authors:** Nazario Bosco, Aleah Goldberg, Adam F Johnson, Xin Zhao, Joseph C Mays, Pan Cheng, Joy J Bianchi, Cecilia Toscani, Lizabeth Katsnelson, Dania Annuar, Sally Mei, Roni E Faitelson, Ilan Y Pesselev, Kareem S Mohamed, Angela Mermerian, Elaine M Camacho-Hernandez, Courtney A Gionco, Julie Manikas, Yi-Shuan Tseng, Zhengxi Sun, Somayeh Fani, Sarah Keegan, Scott M Lippman, David Fenyö, Stefano Santaguida, Teresa Davoli

**Affiliations:** Institute for Systems Genetics and Department of Biochemistry and Molecular Pharmacology, NYU School of Medicine, New York, NY 10016, USA.; Volastra Therapeutics, 1361 Amsterdam Ave Suite 520, New York, NY 10027, USA; Department of Experimental Oncology, IEO European Institute of Oncology IRCCS and Department of Oncology and Hemato-Oncology, University of Milan, Milan, 20141 Italy; Department of Pathology and Laura & Isaac Perlmutter Cancer Center, NYU School of Medicine, New York, NY, USA; Department of Microbiology and Immunology, Weill Cornell Medicine 1300 York Ave, New York, NY; Moores Cancer Center, University of California San Diego, 3855 Health Sciences Drive, La Jolla, CA 92093, USA

**Author notes:** Equal contribution (Nazario Bosco, Aleah Goldberg, Adam F Johnson, Xin Zhao).

## Abstract

Aneuploidy, the presence of chromosome gains or losses, is a hallmark of cancer and congenital syndromes. Here, we describe KaryoCreate (<u>Karyo</u>type <u>CR</u>ISPR <u>E</u>ngineered <u>A</u>neuploidy <u>Te</u>chnology), a system that enables generation of chromosome-specific aneuploidies by co-expression of a sgRNA targeting chromosome-specific CENPA-binding ɑ-satellite repeats together with dCas9 fused to a mutant form of KNL1. We designed unique and highly specific sgRNAs for 19 out of 24 chromosomes. Expression of these sgRNAs with KNL1^Mut^-dCas9 leads to missegregation and induction of gains or losses of the targeted chromosome in cellular progeny with an average efficiency of 8% and 12% for gains and losses, respectively (up to 20%), tested and validated across 9 chromosomes. Using KaryoCreate in colon epithelial cells, we show that chromosome 18q loss, a frequent occurrence in gastrointestinal cancers, promotes resistance to TGFβ, likely due to synergistic hemizygous deletion of multiple genes. Altogether, we describe a novel technology to create and study chromosome missegregation and aneuploidy in the context of cancer and beyond.

**Highlights:**

- We designed sgRNAs targeting chromosome-specific centromeres across 19 human chromosomes
- KaryoCreate combines chromosome-specific centromeric sgRNAs with dCas9 fused to a mutant form of KNL1.
- KaryoCreate allows engineering gains and losses of specific human chromosomes.
- Engineered Chromosome 18q loss promotes tumor-associated phenotypes in colon-derived cells.
- KaryoCreate is a CRISPR-based technology to foster the study of centromere biology and aneuploidy.

## INTRODUCTION

Aneuploidy, the presence of chromosomal gains or losses in a cell, is rare in normal tissues and detrimental for organismal development (Knouse et al., 2014). Some congenital trisomies of specific chromosomes (chr) may be tolerated in humans, such as in Down syndrome (chr 21), Patau syndrome (chr 13), or Edwards syndrome (chr 18), while autosomal monosomies are always lethal (Kirby, 2017). Furthermore, congenital aneuploidies of sex chromosomes such as Klinefelter syndrome (XXY) are more frequent, likely due to the mitigating effect of X-inactivation mediated gene dosage compensation (Groth et al., 2013). The negative consequences of aneuploidy both at the cellular and organismal level are due to a combination of chromosome-specific effects (dependent on the affected chromosome) and chromosome non-specific effects (due to cellular stress phenotypes ascribed to dosage imbalance of many genes, independent of the affected chromosome) (Gordon et al., 2012; Santaguida and Amon, 2015).

Despite its detrimental effect on the physiology of normal cells, aneuploidy is frequent in cancer, estimated to be present in 79% (chromosome level) and 87% (arm level) of solid tumors and 35% (chromosome level) and 40% (arm level) of hematological tumors (Knouse et al., 2017). The number of chromosome or arm gains and losses increases as lesions progress from a premalignant stage, where few (3 chromosome events or 7 arm events on average) are present, to invasive tumors and metastases where many more events are observed (6 chromosome events or 12 arm events on average) (Beroukhim et al., 2010; Davoli et al., 2013; Taylor et al., 2018; William et al., 2021). One interesting feature of aneuploidy in cancer is that it is generally chromosome-specific, i.e. there are specific chromosomes that tend to be gained or lost much more frequently than others (Knouse et al., 2017). We and others recently proposed that these recurrent patterns of aneuploidy are selected for in cancer to maximize the dosage of oncogenes and minimize the dosage of tumor suppressor genes (Davoli et al., 2013; Watkins et al., 2020).

A challenge in studying congenital and cancer aneuploidy is the lack of efficient, straightforward methods for generating aneuploid cell models in which a specific chromosome is added or removed. Widely used methods to induce aneuploidy utilize chemical inhibition, overexpression, or knock down of genes involved in chromosome segregation, such as MPS1, MAD2, or BUBR1, resulting in random chromosome gains or losses (Santaguida and Amon, 2015).

Methods to induce specific chromosome gains are particularly lacking. Microcell-mediated chromosome transfer can be used to transfer specific chromosomes from one donor cell to another, resulting in a specific chromosome gain, but this method is complicated and inefficient, especially in primary cells (Fournier, 1981; Stingele et al., 2012). Engineering of specific chromosome losses has also been limited to few methods and restricted to a subset of chromosomes. These technologies involve the use of CRISPR/Cas9 to eliminate all or part of chromosomes, introducing DNA double strand breaks and potentially instigating undesired genomic instability. One example is the deletion of the short arm of chromosome 3 after insertion of a telomeric sequence in proximity to its centromere (Taylor et al., 2018). Other groups have used CRISPR/Cas9 to cut in non-coding regions to generate whole chromosome losses, but this method has only been successful with the acrocentric chromosomes Y, 14, and 21 (Rayner et al., 2019; Zuo et al., 2017). Finally, additional methods were recently described that use non-centromeric repeats to induce specific losses and, to a lower extent, gains of only chromosomes 1 and 9, with losses occurring mostly at the arm-level (Tovini et al., 2022; Truong et al., 2022).

Human centromeres are composed of highly repetitive AT-rich repeats, also known as α-satellite DNA. Centromeric sequences are hierarchically organized in megabase-long arrays (HOR) of alphoid DNA that directly binds to CENPA, a histone H3 variant, critical to kinetochore position and assembly (Barra and Fachinetti, 2018; Hayden, 2012; Schueler and Sullivan, 2006; Whinn et al., 2019). In the human genome, HOR arrays are made of sequences that are, for the most part, *specific* to individual chromosomes. 15 autosomes and the 2 sex chromosomes have centromeric arrays that are unique to each chromosome (Hayden, 2012). Among the other 7 chromosomes, we can distinguish two families based on centromere similarities: one family comprising chromosomes 1, 5, and 19, and another comprising the acrocentric chromosomes 13, 14, 21, and 22. The centromeres of these chromosomes have been fully resolved (in the cell line CHM13) thanks to recent sequencing efforts by the Telomere-to-Telomere (T2T) consortium (Altemose et al., 2022).

The main function of CENPA-bound centromeric sequences is directing the assembly of the kinetochore, a multiprotein complex that regulates the binding of microtubules to mitotic chromosomes for proper segregation (Musacchio and Desai, 2017). The structure and function of the kinetochore are highly complex and the subject of active investigation. The binding and the regulation of the kinetochore-microtubule attachment are mediated by KMN network (KNL1/MIS12 complex/NDC80 complex), a protein complex located in the outer part of the kinetochore (Cheeseman, 2014). In this complex, the calponin-homology (CH) domains of NDC80 and NUF2 are adjacent and interact directly with microtubules (Korenbaum and Rivero, 2002). During mitosis, each sister kinetochore must be properly attached to one of the opposite poles of the mitotic spindle (amphitelic attachment) to allow their equal segregation into the two daughter cells (Musacchio, 2015). Chromatids that are properly attached experience mechanical tension at the kinetochore, the lack of which activates the spindle assembly checkpoint (SAC), preventing the onset of the metaphase-to-anaphase transition (Musacchio, 2015; Stern and Murray, 2001). SAC triggers the kinase activity of Aurora B, which destabilizes kinetochore-microtubule attachments by phosphorylating several targets in the KMN network, including NDC80 and KNL1. The kinase activity of Aurora B is counteracted by the action of the phosphatase PP1, which is recruited to the kinetochores through direct interaction with KNL1 (Liu et al., 2010). In this highly dynamic process, chromatids under insufficient tension detach from the microtubules so that new attachments can form, until all chromosomes have proper attachment before segregation can occur. The balance between kinase and phosphatase activities ultimately determines the fate of the kinetochore-microtubule attachment and the timing of the metaphase to anaphase transition.

In this work, we describe KaryoCreate, a new method based on the CRISPR/Cas9 technology that combines the chromosome specificity of human centromeric α-satellite repeats with interfering with normal functions of the KMN network (in particular KNL1) to generate chromosome-specific aneuploidy. For 19 out of 24 chromosomes, we computationally predict unique sgRNAs binding >400 times at the centromere with a specificity of 99%. Using KaryoCreate, we demonstrated the successful induction of chromosome-specific aneuploidy for 10 chromosomes tested. In principle, KaryoCreate can be used for 19 out of 24 chromosomes, with the exception of chromosomes sharing similar centromeric sequences such as acrocentric chromosomes. However, we show that induction of gains and losses for the remaining chromosomes is still possible by using sgRNAs targeting both the chromosome of interest and other chromosomes sharing centromeric sgRNA binding sites (instead of single chromosomes). Furthermore, we stably generated two highly recurrent aneuploidies in human gastro-intestinal cancers (chromosome 7 gain and 18q loss), and present data supporting tumor-associated phenotypes associated with chromosome 18q loss in colorectal cells.

## RESULTS

### Computational prediction of sgRNAs targeting chromosome-specific α-satellite centromeric repeats

To design sgRNAs that specifically target the centromere of each chromosome, we used the human genome assembly available from the T2T consortium, the most complete genome assembly available to date (Nurk et al., 2022). To reduce the risk of bias associated with a single assembly, we confirmed that the sgRNA sequences predicted from the T2T assembly (derived from cell line CHM13) could also be found in the hg38 reference genome (Schneider et al., 2017), as there is variation between genome assemblies, including at centromeres (Sullivan and Sullivan, 2020; Willard, 1991).

To maximize the likelihood of interfering with chromosome segregation, we focused our centromeric sgRNA design to target the centromeric HOR that were found to bind to CENPA in ChIP experiments and are thus predicted to bind within the kinetochore assembly region; these HOR are defined as “Live”, or HOR_L, in the T2T assembly (Altemose et al., 2022; Uralsky et al., 2019). For any given chromosome, the ideal sgRNA would 1) have high on-target specificity (i.e. it would not bind to other centromeres or to other locations in the genome), 2) bind to a high number of sites, and 3) have a high efficiency in tethering dCas9 to the genome. For each chromosome, we started by identifying all possible sgRNAs targeting its CENPA-binding α-satellite repeats. We considered the HOR_L arrays and searched for all possible sequences corresponding to the sgRNA target pattern (20 nucleotides followed by NGG PAM). We performed this analysis for all 24 human chromosomes. Next, we determined a set of parameters that define the specificity and efficiency for each candidate sgRNA. For the specificity, we determined the *chromosome specificity* score for each sgRNA, defined as the ratio of the number of binding sites on the centromere of the target chromosome to the total number of binding sites across all centromeres on all chromosomes (multiplied by 100), as well as the *centromere specificity* score, defined as the ratio of the number of binding sites in centromeric regions to the number of sites across the whole genome (multiplied by 100; best score is 100%). We also predicted the efficiency of each sgRNA based on three parameters: GC content (shown to greatly reduce the efficiency of sgRNAs when too high or too low; (Wang et al., 2014)), scores for sgRNA acitvity based on published studies (Doench et al., 2016) (see **Methods**), and total number of binding sites to the specific centromere (Fig. 1A). Finally, as mentioned above, in order not to bias our sgRNA prediction to one assembly, we restricted our search to sgRNAs whose binding site sequence is also found in the hg38 genome assembly.

**Figure 1.**
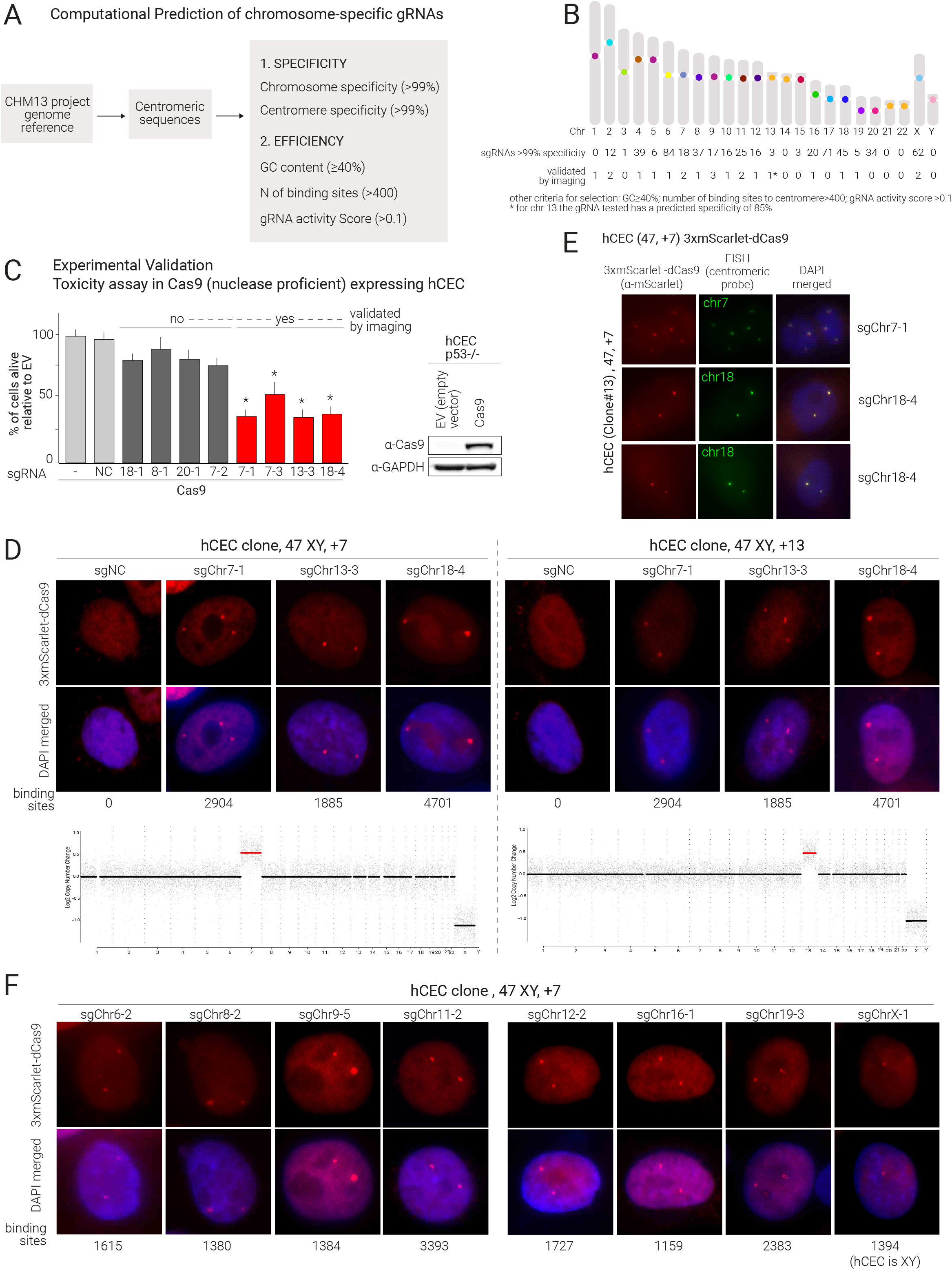
Prediction and validation of chromosome-specific sgRNAs targeting human α-satellite centromeric sequences. (A) Prediction of centromeric sgRNAs specific for each human chromosome based on the CHM13 human reference genome. The two main criteria on specificity and efficiency used for sgRNA selection are reported (see text for details, N=number). (B) Graphic representation of the human karyotype with number of predicted sgRNAs with specificity ≥99% for each human chromosome with the indicated parameters. Number of sgRNAs validated by imaging are also indicated (see panels 1C and 1D, F and Fig. S1). (C) Left Panel: Toxicity assay of centromeric sgRNAs in hCEC hTERT p53-/- expressing Cas9 or empty vector (EV). Nomenclature for sgRNAs: sgRNAα-β refers to a sgRNA that is specific for chromosome α while β refers to the serial number of sgRNAs designed for this chromosome. The percentage of live cells relative to EV was determined 7 days after transduction. Mean and standard deviation are shown; p-values are from Wilcoxon test comparing each condition with NC (*=p<0.05). We also show whether the sgRNAs were (yes) or were not (no) validated by imaging (as in Fig. 1D); the sgRNAs that could not be validated did not show any foci by imaging. Right Panel: Western blot showing Cas9 expression in the hCEC cells used for the toxicity assays. (D) Validation of centromere targeting in hCEC hTERT p53-/- clones (containing 3 copies of chr7 or chr13 respectively) expressing 3xmScarlet-dCas9 fusion. Representative images of interphase cells with the indicated sgRNAs and percentage of cells displaying the foci. The number of binding sites for each sgRNA is also indicated. Low-pass WGS is also shown to confirm specific aneuploidies in the two clones. (E) IF/FISH using chromosome 7 and 18 specific probes indicating colocalization with 3xmScarlet-dCas9 fusion (as in D) in hCEC hTERT p53-/- cells (trisomic for chr7) co-expressing sgRNA7-1 or sgRNA18-4. FISH-IF was performed using anti-mScarlet antibody (red) and chromosome 7/18 centromeric FISH probes (green). (F) Additional sgRNAs validation in hCEC hTERT p53-/- cells (clone 13) expressing the fluorescently-tagged dCas9 (3xmScarlet-dCas9 fusion) for the indicated sgRNAs. Representative images of interphase cells are shown as well as the % of cells displaying the foci.

Using a threshold of 99% for both *chromosome specificity* score and *centromere specificity* score, a GC content >40%, a minimum of 400 sgRNA binding sites, an *activity score* (Doench et al., 2016) >0.1 and representation in hg38, we designed at least one specific sgRNA for 19 out of 24 chromosomes (Fig. 1B). Among the predicted sgRNAs using these criteria, the median and mean predicted number of binding sites were 1376 and 1590. Increasing the chromosome specificity score from 99% to 100% resulted in at least one sgRNA for 16 out of 24 chromosomes, highlighting the specificity of our approach. The remaining 5 chromosomes (1, 14, 21, 22, & Y) can be targeted in pairs or small groups using a sgRNA specific to their shared centromere sequences (see below, Fig. 4B).

### Experimental validation of sgRNAs targeting α-satellite centromeric repeats on 15 human chromosomes

To experimentally validate the predicted sgRNAs in human cells, we expressed each sgRNAs in the presence of endonuclease proficient Cas9 and performed a toxicity assay based on the prediction that several double-strand breaks at the centromere are likely to decrease cell viability. We used hTERT TP53-KO human colonic epithelial cells (hCEC, (Ly et al., 2011)) and hTERT retinal pigment epithelial cells (RPE) expressing p21 and RB shRNAs (Maciejowski et al., 2015). We transduced Cas9-expressing RPEs and hCECs with a lentiviral vector (pLentiGuide-Puro-FE, see **Methods**) expressing either a negative control sgRNA (sgNC; (Sanjana et al., 2014)) that does not target any sequence in the human genome or centromeric sgRNAs. After selection, we plated the same number of cells for each sample and counted the total number of cells after 5-6 days. Hereafter we use the following nomenclature for sgRNAs: sgChrα-β where α refers to the chromosome that the sgRNA is specific for and β refers to the serial number of sgRNA designed for this chromosome. We initially tested sgRNAs predicted for chromosomes (chr) 7 (sgChr7-1, sgChr7-2, sgChr7-3), 13 (sgChr13-1, sgChr13-2, sgChr13-3), and 18 (sgChr18-1, sgChr18-2, sgChr18-3, sgChr18-4). Compared to controls (sgNC or non-transduced cells), hCEC and RPE expressing sgChr7-1, sgChr7-3, sgChr13-3, or sgChr18-4 exhibited at least 50% reduction in cell number, while the other sgRNAs did not result in a significant difference in viability (Fig. 1C; **Fig. S1A**). We selected the most toxic sgRNA for each chromosome for additional testing.

To confirm that the sgRNAs were indeed targeting the intended centromeres, we designed a dCas9-based imaging system consisting of 3 mScarlet fluorescent molecules fused to the N-terminus of endonuclease dead Cas9 (3xmScarlet-dCas9). To achieve a consistently high level of fluorescent construct expression, we FACS-sorted 3xmScarlet-dCas9-transduced hCECs for high levels of mScarlet-derived fluorescent signal. As we expected, 3xmScarlet-dCas9-transduced hCECs cells expressing sgChr7-1, sgChr13-3, or sgChr18-4 showed the formation of bright foci in their nuclei (Fig. 1D). Notably, the sgRNAs that did not exhibit toxicity in the presence of Cas9 failed to form foci in the nucleus visible by microscopy (Fig. 1C and data not shown).

To further confirm the chromosome-specificity of the sgRNA, we used two independent approaches. We first used hCEC clones with known aneuploidies to verify whether the observed number of foci was consistent with the expected number based on whole genome sequencing (WGS)-based copy number analysis. We found that hCEC lines containing 3 copies of chromosome 7 or 13 showed three foci, when transduced with sgChr7-1 or sgChr13-3, respectively (Fig. 1D). As anticipated, transduction with sgRNAs targeting disomic chromosomes lead to the formation of two foci per nucleus (Fig. 1D). We note that the percentage of cells showing foci was 45% in the polyclonal population of hCEC transduced with 3xmScarlet-dCas9 and increased to 72% in a monoclonal population expressing high levels of 3xmScarlet-dCas9 derived from the hCEC clone trisomic for chromosome 7 (**Fig. S1B**). The increased expression in 3xmScarlet-dCas9 in the monoclonal population likely explains this result and may have implications for the efficiency of KaryoCreate (see below and Discussion). Next, we confirmed that the sgRNA-mediated 3xmScarlet-dCas9 foci were indeed localizing at specific centromeric sequences by Fluorescence In-Situ Hybridization (FISH) using centromeric probes. We found colocalization of mScarlet foci with FISH signals for both chromosome 7 (sgChr7-1) and 18 (sgChr18-4) (Fig. 1E). Altogether, these experiments support the fact that the computationally predicted sgRNAs are capable of recruiting dCas9 to the expected specific centromere.

We experimentally tested the capability to induce foci for additional sgRNAs in hCECs or RPEs (**Fig. S1C-D**). We expressed 75 individual sgRNAs in hCECs individually with3xmScarlet-dCas9 and confirmed the formation of the expected number of foci for 24 sgRNAs targeting in a specific manner 15 different chromosomes, namely chromosomes 2, 4, 5, 6, 7, 8, 9, 10, 11, 12, 13, 16, 18, 19, X (Fig. 1F, **Fig. S1C**). We also confirmed 4 sgRNAs targeting 4 different chromosomes in RPE cells (**Fig. S1D**). Given the fact that RPEs and hCECs are from different individuals, and both supported the validation of sgRNAs (predicted based on CHM13 cells and hg38), this suggests that the sgRNAs are likely to function across different cell lines.

In summary, the average sgRNA validation rate was 32% but varied widely among chromosomes. For most chromosomes, we were able to find a good sgRNA among the top 3 predicted ones. For other chromosomes (e.g. 20) we were not able to validate a unique sgRNA among 8 tested, potentially due to the fact that the total number of binding sites of these sgRNAs to the centromere is generally lower than for other chromosomes.. The predicted sgRNA efficiency score did not correlate with the ability of the sgRNA to form foci (**Fig. S1E**, left panel). Instead, for the sgRNAs that formed foci, there was a significant positive correlation between the intensity of the signal of the foci and the number of binding sites at the centromeres predicted based on the CHM13 genome reference (r=0.65, p=0.03; **Fig. S1E**, right panel).

Altogether, we have designed and validated 24 chromosome-specific sgRNAs targeting the centromeres of 15 different human chromosomes.

### Centromeric targeting of KNL1^Mut^-dCas9 induces modest mitotic delay, chromosome missegregation and micronuclei formation

To induce chromosome missegregation, we built and tested four fusion partners for dCas9 that we predicted would disrupt proper kinetochore-microtubule attachments (Fig. 2A, **Fig. S2A**). Two constructs (KNL1^S24A;S60A^-dCas9 and KNL1^RVSF/AAAA^-dCas9) utilize the N-terminal portion (amino acid (aa) 1-86) of KNL1, containing mutations with opposing effects, both predicted to disrupt the cross-regulation between Aurora B and PP1 (**Fig. S2A**). KNL1^S24A;S60A^ is predicted to be always bound to PP1 as its mutated residues cannot be phosphorylated by Aurora B, which would normally inhibit the interaction of KNL1 with PP1 (**Fig. S2A**); thus this mutant is predicted to disrupt Aurora B-mediated regulation by aberrantly recruiting PP1 to kinetochores. On the other hand, KNL1^RVSF/AAAA^ (Liu et al., 2010) contains a mutation of the RVSF motif (aa 58-61) which cannot interact and recruit PP1 to the centromere (**Fig. S2A**); this mutant is predicted to alter the normal error correction mediated by the cross-regulation between Aurora B and PP1. The other two constructs (NDC80-CH1-dCas9 and NDC80-CH2-dCas9) were designed to render the interaction between kinetochores and microtubules hyperstable and unamenable to destabilization by Aurora B. These constructs contain one (NDC80-CH1) or two (NDC80-CH2) mutant CH domains of NDC80 (aa 1-207), the region of NDC80 that, together with NUF2, binds to microtubules. These CH domains contain 6 residues whose phosphorylation by Aurora B inhibits the interaction with microtubules (DeLuca et al., 2006); these 6 residues are all mutated in our constructs (DeLuca et al., 2006) thus preventing Aurora B-mediated regulation (**Fig. S2A**). Western blot analysis of transduced cells showed that KNL1^RVSF/AAAA^-dCas9 and KNL1^S24A;S60A^-dCas9 expression levels were higher than NDC80-CH1-dCas9 and NDC80-CH2-dCas9 (**Fig. S2B**). For the KNL1 constructs, the N-terminal fusions were generally more stable than the C-terminal fusions (**Fig. S2C**, Fig. 2A).

**Figure 2.**
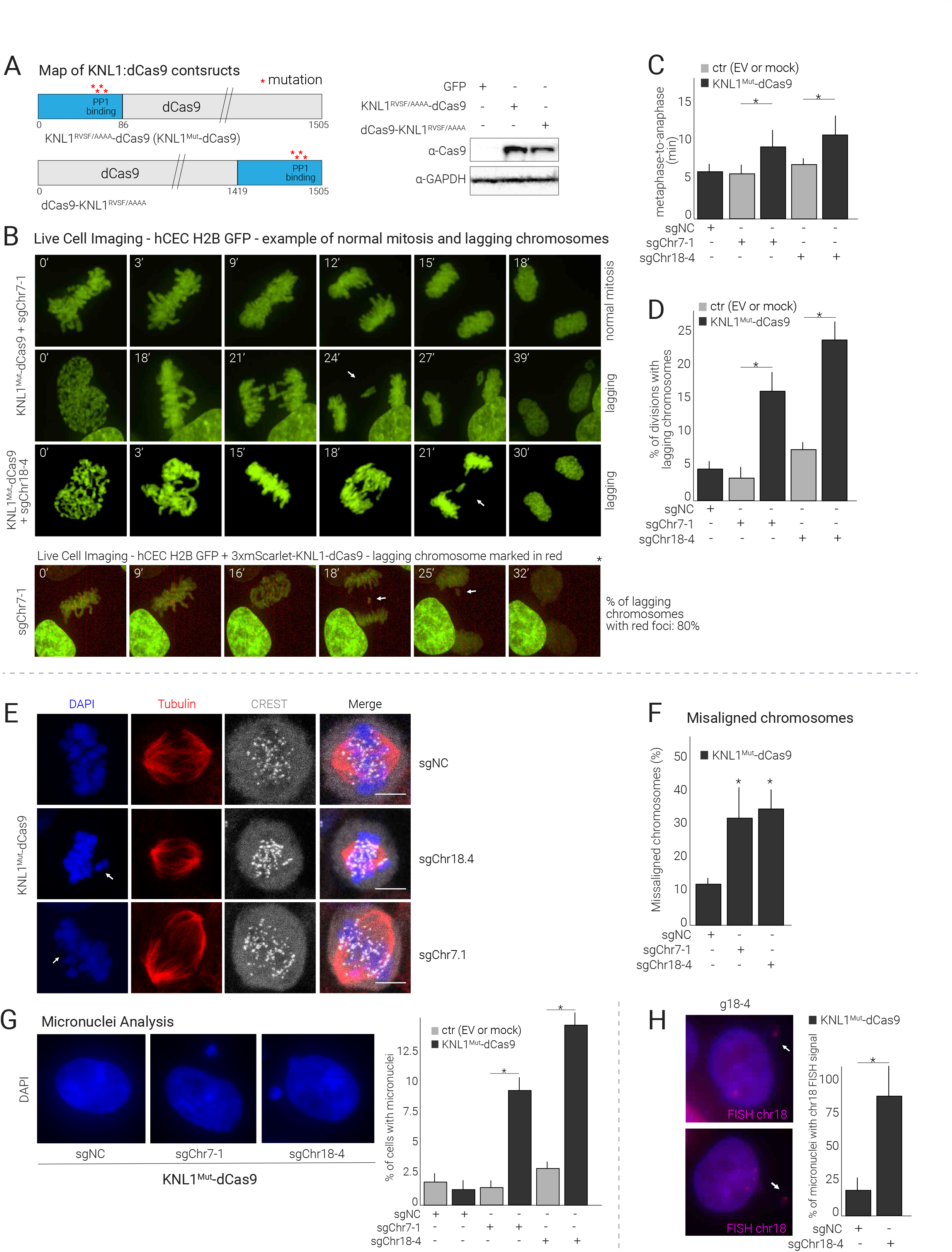
KNL1^Mut^-dCas9 targeted to centromeres induces modest mitotic delay and chromosome missegregation. (A) **Left**: Map of constructs used to test KaryoCreate: KNL1^RVSF/AAAA^-dCas9 and dCas9-KNL1^RVSF/AAAA^. See text for details. **Right**: Western blot showing the expression of the indicated construct in hCEC cells. (B) Time-lapse imaging of hCEC hTERT p53-/- cells expressing H2B-GFP, KNL1^Mut^-dCas9 and sgChr7-1 or sgChr18-4 or NC as control. Cells were analyzed for time spent in mitosis and for the presence of lagging chromosomes (quantified in C and D respectively). In the lower panel, the analysis was performed in hCEC hTERT p53-/- H2B-GFP cells co-expressing 3xmScarlet-KNL1^Mut^-dCas9 and sgChr7-1, indicating specific chromosome missegregation. Experiments performed in duplicates or triplicates and at least 20 cells dividing were analyzed for each condition. Complete time-lapse series are included as Movies S1-S4. (C) Quantification of mitotic duration calculated as time spent between metaphase and the onset of anaphase from analysis in Fig. 2B. Mean and standard deviation are shown; p-values are from Wilcoxon test. (D) Quantification of mitoses showing lagging chromosomes from the imaging performed in Fig. 2B. Mean and standard deviation are shown; p-values are from Wilcoxon test. (E) IF showing congression defects in HCT116 cells expressing KNL1^Mut^-dCas9 and sgChr7-1 or sgChr18-4. Spindle is stained with tubulin (red), DNA with DAPI (blue), and CREST (gray) to visualize aligned and misaligned kinetochores. (F) Quantification of chromosome congression defects for the experiment described in Fig. 2E. Mean and standard deviation from triplicates are shown. (G) Micronulclei analysis in hCEC hTERT p53-/- expressing KNL1^Mut^-dCas9 and with sgChr18-4, 7-1, and NC. Cellswere fixed at 7-10 days after transduction with the specific sgRNAs. Quantification on the right. The percentage of live cells relative to EV was determined 7 days after transduction. Mean and standard deviation from triplicates are shown; p-values are from Wilcoxon test (*=p<0.05). (H) FISH indicating Chr18-containing micronuclei in hCEC hTERT p53-/- cells (treated as in 2G), quantification of micronuclei counts is shown; experiment performed in duplicates.

Given the higher protein expression level and the increased efficiency in inducing chromosome gains and losses compared to the other constructs (see next section and **Fig. S3B**), we focused on the KNL1 constructs, particularly KNL1^RVSF/AAAA^-dCas9, hereafter referred to as KNL1^Mut^-dCas9. To confirm centromeric localization of the fusion protein, we transduced hCECs expressing a fluorescently tagged version of the KNL1^Mut^-dCas9 (3xmScarlet-KNL1^Mut^-dCas9) with different sgRNAs. We observed the expected number of foci in the presence of different centromeric sgRNAs (e.g. sgChr7-1 and sgChr18-4) (**Fig. S2D**), indicating that the fusion of KNL1^Mut^ with dCas9 does not alter the ability of dCas9 to be recruited to centromeres. Next, using live-cell imaging, we examined the effect of KNL1^Mut^-dCas9 on the duration of mitosis and chromosome segregation. hCEC constitutively expressing GFP-tagged histone H2B were transduced with KNL1^Mut^-dCas9 or empty vector (EV) and with sgChr7-1, sgChr18-4 or sgNC. Cells expressing KNL1^Mut^-dCas9 and either sgChr7-1 or sgChr18-4 showed a slower progression through mitosis compared to cells transduced with EV and either sgChr7-1 or sgChr18-4 (Fig. 2C): the average time spent in the metaphase to anaphase transition increased from 6 minutes to 9 or 10 minutes in the sgChr7-1 or sgChr18-4 condition, respectively (Fig. 2B, C). Even though this phase of mitosis was about 50% slower, cells did not arrest in metaphase but completed mitosis. Consistently, there was only a slight and not significant decrease in the proliferation rate compared to the sgNC (**Fig. S2E**). The number of cell divisions with lagging chromosomes increased from less than 5% to 15% between EV + sgChr7-1 and KNL1^Mut^-dCas9 + sgChr7-1 and from 7% to 23% between EV + sgChr18-4 and KNL1^Mut^-dCas9 + sgChr18-4 (Fig. 2B, D). Furthermore, live-cell imaging of cells expressing 3xmScarlet-KNL1^Mut^-dCas9 and sgChr7-1, where mScarlet marks chromosome 7 as in **Fig. S2D** (polyclonal population), showed that about 80% of the lagging chromosomes observed during mitosis showed red foci, supporting that the missegregation was chromosome-specific (Fig. 2B, lower panel); unfortunately in this specific type of experiment, it was not possible to use the sgNC as a control as no foci can be formed.

To corroborate these data in a different cell line, we performed a similar experiment in the HCT116 (TP53 WT) colon cancer cell line. HCT116 cells were transduced with KNL1^Mut^-dCas9 and sgNC, sgChr7-1 or sgChr18-4. The cells were fixed and stained with fluorescent antibodies against Tubulin to visualize the mitotic spindle, with CREST serum to visualize the centromere and counterstained with DAPI in order to assess chromosome alignment in mitosis (Fig. 2E). Consistent with previous results in hCEC TP53 KO, the fraction of mitoses with misaligned chromosomes increased from 12% in the sgNC samples to 32% and 35% in the sgChr7-1 and sgChr18-4 conditions, respectively (Fig. 2F).

Finally, since a well-known readout of missegregation is the formation of micronuclei, we scored the fraction of KNL1^Mut^-dCas9-expressing hCECs containing micronuclei 7-10 days after transduction with sgNC, sgChr7-1, or sgChr18-4. The percentage of cells showing micronuclei increased from less than 2.5% for sgNC to 9% for sgChr7-1 and 14% for sgChr18-4 (Fig. 2G). Furthermore, we performed FISH using a chr 18 centromeric probe on cells expressing KNL1^Mut^-dCas9 and sgChr18-4 and found that 85% of micronuclei showed a FISH signal, indicating that more than 75% of micronuclei contained chromosome 18. Altogether, these data indicate that the tethering of KNL1^Mut^-dCas9 with centromere-specific sgRNAs can induce chromosome misalignment, lagging chromosomes, modest mitotic delay and micronuclei formation of the targeted chromosome without major effects on the rate of cell division.

### KaryoCreate allows induction of chromosome-specific gains and losses in human cells

Having designed and validated chromosome-specific sgRNAs and dCas9-based constructs for induction of chromosome missegregation, we next tested the capability of this system, designated “KaryoCreate” for <u>Karyo</u>type <u>CR</u>ISPR <u>E</u>ngineered <u>A</u>neuploidy <u>Te</u>chnology, to generate human cell lines with chromosome-specific gains and losses (Fig. 3A). We reasoned that the ideal system would be based on a transient expression of the constructs to allow subsequent generation of stable cell lines containing the aneuploidy of interest. Therefore, we designed a system allowing doxycycline-inducible expression of KNL1^Mut^-dCas9 using pIND20 vector (Meerbrey et al., 2011), and a constitutive lentiviral vector pLentiGuide-Puro-FE for the sgRNA expression (Fig. 3B, 3C, see **Methods**).

**Figure 3.**
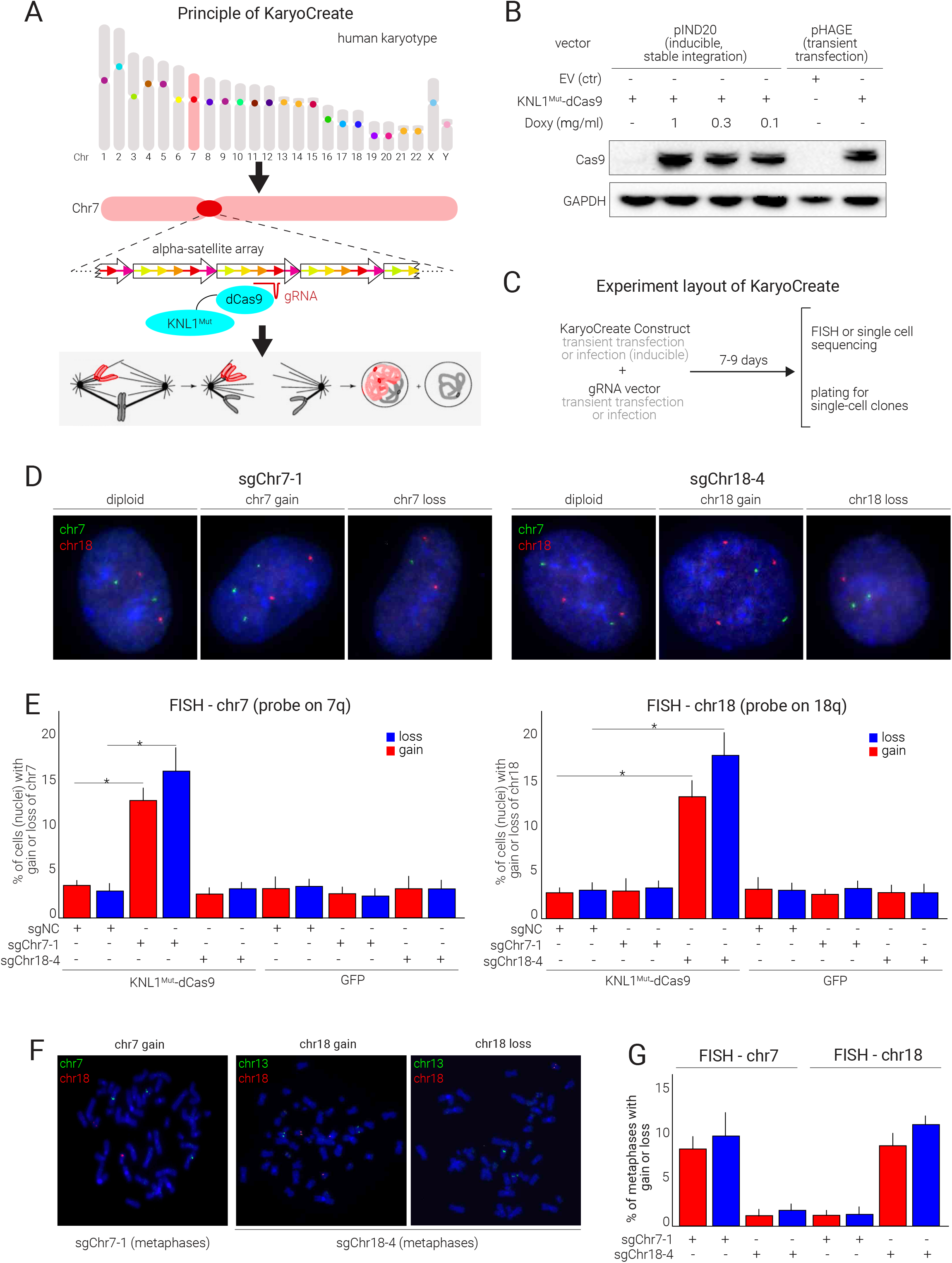
KNL1^Mut^-dCas9 targeted to centromeres allows induction of chromosome-specific gains and losses in human cells. (A) Overview of KaryoCreate method: the chromosome-specificity of human α-satellite centromere sequences allow targeted missegregation while leaving the other chromosomes unaffected. KaryoCreate combines the sequence specificity of the sgRNA/CRISPR system to tether to the specific centromere a fusion protein that induce dysfunctional kinetochore-microtubule interactions and subsequent aneuploidy (KNL1^Mut^ fused to dCas9 represented in the figure, see text for details). See text and Fig. S2 for additional details. (B) Western blot showing the expression of the indicated construct in hCEC cells, either using transient transfection with a constitutive promoter (pHAGE-CMV) or using lentiviral infection with a doxycycline-inducible promoter (pIND20). Doxy: doxycycline. (C) Schematics showing the timeline of a KaryoCreate experiment involving transient expression of the chromosome-specific sgRNA and of KNL1^Mut^-dCas9; generally cells are harvested after 7 to 9 days and then analyzed by FISH or single cell sequencing. Cells can eventually be plated to derive single cell clones. (D) FISH using probes specific for chr7 and chr18 on hCEC hTERT p53-/- diploid cells 10 days after expression of KNL1^Mut^-dCas9 and an sgRNA (negative control, sgChr7-1 or sgChr18-4; see also Fig. 1E). Representative images of control cells (no gains/losses: 2 foci for chr7 and 2 foci for chr18) and cells showing gains or losses of chr7 or 18 are shown. (E) FISH quantification for chr7 (left panel) or chr18 (right panel) for hCEC hTERT p53-/- diploid cells treated as in Fig. 3D (manual quantification). Note that compared to sgNC, expression of sgChr7-1 leads to gains and losses of chr7 but not of chr18 while sgChr18-4 leads to gains and losses of chr18 but not of chr7. Mean and standard deviation are shown among triplicates; p-values are from Wilcoxon test (*=p<0.05). (F) Metaphase spreads from hCEC hTERT p53-/- cells treated as in (3D) and processed by FISH using probes specific for chr7, chr13, and chr18 as indicated. Representative pictures are shown. (G) Quantification of FISH signals from Fig. 3F.

To test KaryoCreate, hCECs were transduced with pIND20-KNL1^Mut^-dCas9 or pIND20-GFP as control and with the vector expressing sgRNAs sgNC, sgChr7-1 or sgChr18-4. Cells were treated with doxycycline for 7-8 days and then analyzed by FISH using probes for chromosomes 7 and 18. As expected, 95% of the control cells (GFP and sgNC) showed two copies of chromosomes 7 and 18 (Fig 3D, E). This percentage did not significantly change in cells expressing the KNL1^Mut^-dCas9 construct and sgNC, indicating that in the absence of a centromere-specific sgRNA, KNL1^Mut^-dCas9 does not induce chromosome gains and losses (Fig. 3D, E). Compared to sgNC, hCECs expressing KNL1^Mut^-dCas9 with sgChr7-1 showed a significant increase in the percentage of cells showing 1 copy (from 3 to 16%; p=0.01) or >2 copies (from 2.8 to 12.5%; p=0.03) of chromosome 7, while the percentage of cells showing 1 or 3 copies of chromosome 18 was not significantly changed (from 3 to 3.2%). We next tested sgChr18-4, and observed a significant increase in losses (from 2 to 17.5%; p=0.01) and gains (from 2.5 to 14%; p=0.02) of chromosome 18, but not chromosome 7 (Fig. 3D, E). Furthermore, we obtained comparable results when we restricted the FISH analysis to metaphase spreads as opposed to nuclei (Fig. 3F, G). This further supports that cells containing chromosome-specific gains and losses can progress through mitosis as previously shown and suggests that karyotypic changes can be maintained over cell divisions, eventually allowing isolation of clonal aneuploid populations. Generally, the percentage of losses was higher than the percentage of gains (see below and Discussion).

We next developed two additional systems for KaryoCreate. The first system involves transient co-transfection of a CMV-driven KNL1^Mut^-dCas9 (pHAGE vector, not inducible) with a vector expressing the sgRNAs. The second system involves stably integrating both the sgRNA vector and a pHAGE vector containing KNL1^Mut^-dCas9 fused to an FKBP-based degradation domain (Banaszynski et al., 2006) which is stabilized after treatment with the small molecule Shield-1 (see **Methods**). Overall, the three methods of transient KNL1^Mut^-dCas9 expression gave very similar results (**Fig. S3A**).

We next analyzed the frequency of aneuploidy induced by other constructs generated for KaryoCreate (NDC80-CH1-dCas9 and NDC80-CH2-dCas9, described above, see **Fig. S2A-C**). Compared to KNL1^Mut^-dCas9, the other fusion proteins induced aneuploidy with a lower efficiency (**Fig. S3B**). KNL1^S24A;S60A^-dCas9 had similar levels of induced aneuploidy as KNL1^Mut^-dCas9 (KNL1^RVSF/AAAA^-dCas9), while NDC80-CH1-dCas9 and NDC80-CH2-dCas9 constructs showed lower but still appreciable efficiency (see **Fig. S2B, S2C**).

Considering that the expression levels of the different dCas9 fusion constructs correlated with their efficiency of aneuploidy induction, we set out to evaluate which other parameters and conditions could affect KaryoCreate’s efficiency. We found that higher levels of KNL1^Mut^-dCas9 expression induced a higher percentage of aneuploidy: a 3-fold increase in KNL1^Mut^-dCas9 expression led to a two-fold increase in gains or losses (**Fig. S3C-D**). Next, we found that combining multiple sgRNAs targeting the same chromosome (sgChr7-1 + sgChr7-3 or sgChr9-3 + sgChr9-5) did not increase the percentage of aneuploid cells as compared to individual sgRNAs, despite the higher number of predicted binding sites achieved by combining the sgRNAs (**Fig. S3E-F**). Finally, we tested whether FACS sorting, based on a cell surface marker encoded on the target chromosome, could increase the percentage of cells with gains or losses. We identified EPHB4, an ephrin receptor encoded by a gene on chromosome 7 and sorted cells after KNL1^Mut^-dCas9 and sgChr7-1 expression. The percentage of cells with chr 7 gain increased from 12% to 26% from unsorted to high-EPHB4 sorted cells (**Fig. S3G**) and the percentage of cells with chr 7 loss increased from 8% to 16% from unsorted to low-EPHB4 sorted cells. Altogether, the results shown in this section indicate that KaryoCreate is capable of inducing efficient chromosome-specific aneuploidy for the chromosomes tested.

### KaryoCreate allows induction of both arm-level and chromosome-level gains and losses across human chromosomes

Our FISH analyses showed that targeting chromosome 7 does not affect chromosome 18 and vice-versa, but does not rule out erroneous targeting of other chromosomes. To extend our analysis of KaryoCreate’s specificity across all chromosomes, we performed high-throughput single-cell RNA sequencing (scRNAseq) to estimate genome-wide DNA copy number profiles across thousands of KaryoCreate-treated cells (Gao et al., 2021; Patel et al., 2014; Tirosh et al., 2016). To infer copy number by scRNAseq, we developed a modified version of CopyKat (see **Methods**). Briefly, this method uses the median expression of genes across each chromosome arm as an estimate of DNA copy number. The percentage of gains and losses for each arm is then estimated by comparing the DNA copy number distribution of each experimental sample to that of the control population (e.g. aneuploid versus diploid cell lines). To prove the ability to infer the arm-level copy number across chromosomes, we compared scRNAseq across single cells and bulk shallow WGS of two hCEC-derived clonal cell lines that specifically contained chromosome 7 and 19p gains or chromosome 18 losses, and found that patterns of aneuploidy inferred by scRNAseq recapitulated the patterns shown by bulk WGS (**Fig. S4A**). Analysis of a trisomic chromosome 7 clone showed that the percentage of cells with chromosome 7 gain was 91% by FISH and 80% by scRNAseq. Similarly, analysis of a more complex karyotype (+chr7, -chr18, +19p) showed that the percentage of cells with chromosome 7 gain was 88% by FISH and 76% by scRNAseq, and chromosome 18 loss was 87% by FISH and 81% by scRNAseq (**Fig. S4B**). Notably, scRNAseq slightly underestimated the presence of aneuploidy, especially of gains, likely since 2 to 3 copies is an increase in DNA and RNA of 33%, while loss of 1 copy from 2 copies corresponds to a decrease of 50%. Overall, we concluded that scRNAseq is a bona fide method for the analysis of genome-wide gains and losses in single cells.

We performed scRNAseq on diploid TP53-/- hCEC cells 7 days after KaryoCreate for chromosome 7 (sgChr7-1), chromosome 18 (sgChr18-4) and sgNC to estimate the overall frequency of chromosomal gains and losses (Fig. 4A). Using the pIND20 vector to mediate the expression of KNL1^Mut^-dCas9 (Fig. 3B; expression level intermediate compared to the ones reported in Fig. S3C), we successfully detected arm-level copy number variants (CNVs) for most chromosomes, except for chromosomes with zero or a low (<20) number of detected genes on the *p* arm. First, we confirmed that the expression of the KNL1^Mut^-dCas9 with the sgNC construct did not significantly induce aneuploidy compared to an EV control (Fig. 4B, **Fig. S4C**). Notably, the percentage of gains and losses observed across chromosomes for the control (sgNC, Fig. 4B) was very low, with an average of 0.9% for gains and 1.2% for losses. Moreover, we confirmed the induction of chromosome-specific gains or losses after expression of KNL1^Mut^-dCas9 and chromosome-specific sgRNAs, consistent with our FISH experiments (Fig. 3D, E). For example, for chromosome 18 (sgChr18-4), we estimated chromosome 18 in 10% of cells and losses in 17% of cells by scRNAseq (Fig. 4B) and for chromosome 7 (sgChr7-1), we estimated chromosome 7 gains in 9% of cells and losses in 11% of cells by scRNAseq (measured for 7q as for the FISH; Fig. 4B). Most importantly, scRNAseq demonstrated that aneuploidy induction was highly chromosome-specific: the average percentage of gains or losses for all the not targeted chromosomes was on average of 1% (Fig. 4B). It is important to consider that the level of losses and (especially) gains observed in the sgNC across chromosomes (0.9 for gains and 1.2% for losses) level is about 3 times lower than the one observed by DNA FISH (3% for both gains and losses) (Fig. 3E), potentially contributing to the lower percentage of aneuploidy observed by scRNAseq than by FISH in the targeted chromosome (especially for gains).

**Figure 4.**
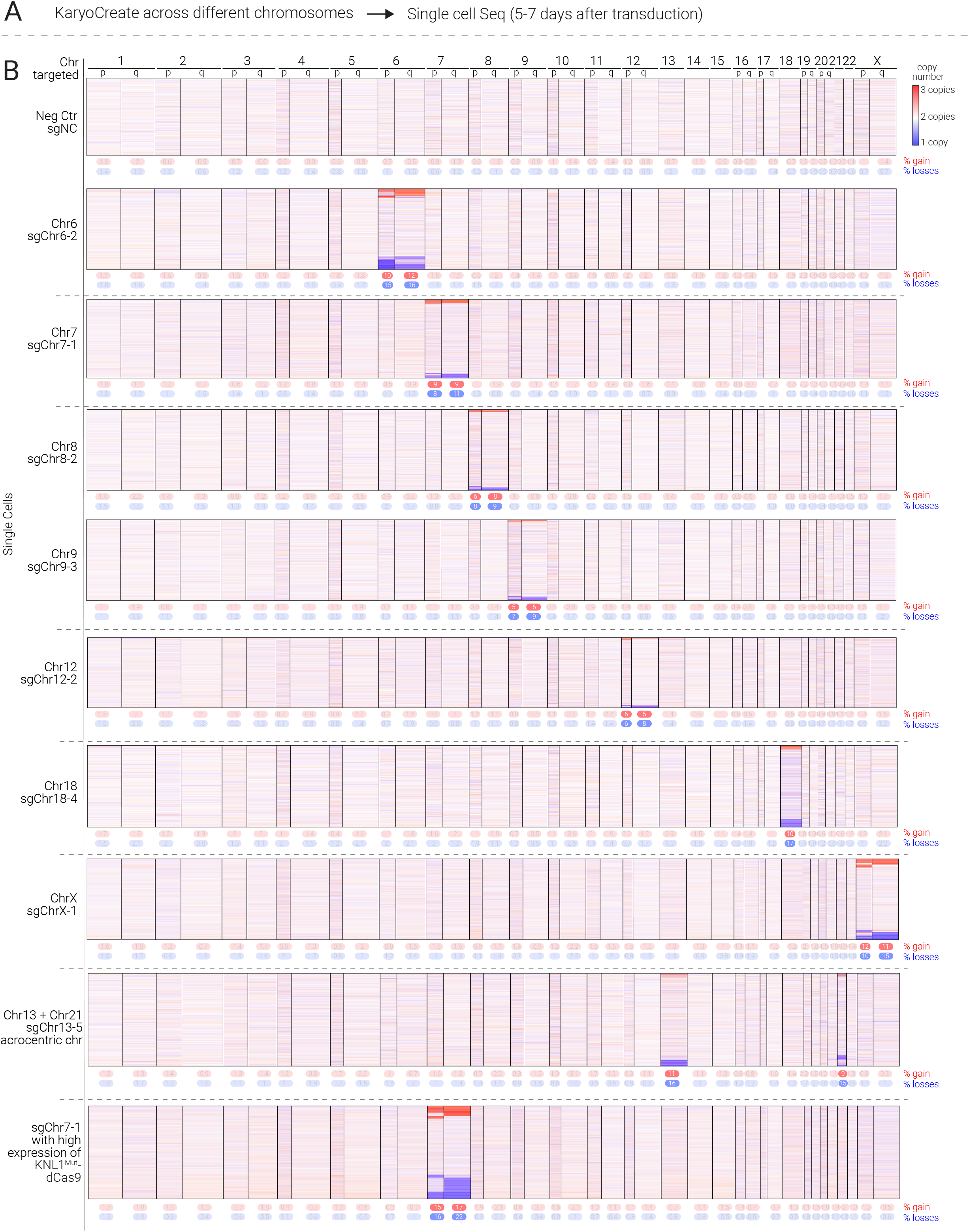
KaryoCreate allows induction of both arm-level and chromosome-level gains and losses across human chromosomes. (A) Schematic of the scRNAseq experiment to validate sgRNAs in hCEC diploid cells. (B) Heatmap depicting gene copy numbers inferred from scRNAseq analysis following KaryoCreate as described in 4A. Individual chromosomes (or combination of chromosomes) were specifically missegregated using KaryoCreate scRNAseq was used to quantify the presence of chromosome or arm-level gains or losses using a modified version of CopyKat. The median expression of genes across each chromosome arm is considered to estimate the DNA copy number. The % of gains and losses for each arm (reported below each heatmap) is estimated by comparing the DNA copy number distribution of each experimental sample (chromosome-specific sgRNA) to the negative control (sgRNA NC; see also **Methods**). In the heatmaps the rows represent individual cells, the columns represent different chromosomes, and the color represents the copy number change (gain in red and loss in blue). In the last sample, ‘higher expression of KNL1^Mut^-dCas9’ indicates that the cells were transduced with a higher amount of the construct (see Fig. S3C).

We further tested KaryoCreate using sgRNAs targeting additional chromosomes, including 6, 8, 9, 12, 16 and X, that were previously validated to induce foci with mScarlet-dCas9 (Fig. 4B, see also Fig. 1 and **Fig. S1**). We performed KaryoCreate with the diploid TP53-/- hCEC cells expressing KNL1^Mut^-dCas9 (pIND20) and analyzed the cells through scRNAseq seven days after transduction with the sgRNAs. In all cases tested, we found that cells expressing the chromosome-specific sgRNAs showed an increase in the percentage of gains and losses in the targeted chromosome compared to the sgNC control. The percentage of chromosome-specific gains and losses differed among the chromosomes and ranged between 5 and 12% for gains (average: 8%) and between 7 and 17% for losses (average: 12%) (Fig. 4B). Importantly, gains or losses of the non-targeted chromosomes were never observed in percentages greater than in the sgNC control.

Furthermore, in agreement with our previous findings, we observed that the expression level of the KNL1^Mut^-dCas9 construct correlated with the efficiency of KaryoCreate in inducing chromosome-specific aneuploidy. In fact, a 3-fold increase in the expression level of KNL1^Mut^-dCas9 (see **Fig. S3C** for WB) resulted in approximately 50% increase in gains from 9% to 16% and losses from 11% to 22% (measured for 7q as for the FISH, Fig. 4B, compare sgChr7-1 and sgChr7-1 with high KNL1^Mut^-dCas9 expression). Finally, we obtained similar results using KaryoCreate in RPE cells using pHAGE-mediated expression of KNL1^Mut^-dCas9 (**Fig. S4D**), suggesting that the method can be applied to different cell lines.

Throughout the scRNAseq analysis, we noted that in addition to whole-chromosome gains and losses, KaryoCreate also induced arm-level gains and losses, where only one of the two chromosomal arms (*p* or *q*) is gained or lost. Across the chromosomes tested, approximately 60% of the aneuploidy events were arm-level events and 40% were whole chromosome events (**Fig. S4E**). On average, there were 28% whole chromosome losses, 17% whole chromosome gains, 32% arm-level gains and 23% arm-level losses (**Fig. S4E**). Thus, KaryoCreate is highly specific for the target chromosome, and induces both chromosome-level and arm-level gains and losses, with a higher proportion of losses (see Discussion).

### Induction of aneuploidy simultaneously across multiple chromosomes and in acrocentric chromosomes

One potential challenge of KaryoCreate is targeting individual chromosomes that have significant centromeric similarity to others. In humans, the acrocentric chromosomes 13, 14, 15, 21 and 22 share a high degree of centromeric sequence similarity (Rayner et al., 2019; Zuo et al., 2017), and thus it is not possible to design chromosome-specific sgRNAs. Given the relevance of these chromosomes in congenital aneuploidies, we asked whether generation of cells containing specific aneuploidies would be possible by using sgRNAs targeting both the chromosome of interest and other chromosomes sharing centromeric sgRNA binding sites (instead of single chromosomes); clones could then be isolated and screened to contain the karyotype of interest. For example, while we could not design sgRNAs targeting chromosome 21 exclusively, we were able to design one sgRNA targeting both chromosome 21 and chromosome 13 (sgChr13-5). We found that expression of KNL1^Mut^-dCas9 in hCEC with sgChr13-5, indeed generated 11% and 16% of cells with chromosome 13 gains or losses, respectively and of 9% and 15% of cells with chromosome 21 gains or losses, respectively (Fig. 4B). Only 5% of cells contained changes in both chromosomes.

Finally, we explored the possibility of generating gains and losses of different chromosomes within the same cell by combining pairs of centromeric sgRNAs. We transduced hCEC with KNL1^Mut^-dCas9 together with sgChr7-1 and sgChr18-4 and found a significant induction of gains and losses for both chromosomes, comparable to the levels observed using the single sgRNAs; furthermore, 8% of the cells had changes in both chromosome 7 and chromosome 18 (**Fig. S4F**). Altogether these data show that KaryoCreate can generate chromosome gains and losses across several individual human chromosomes including acrocentric chromosomes as well as combinations of chromosomes.

### 18q loss in colon cells promotes resistance to TGFβ signaling likely due to haploinsufficiency of multiple genes

As mentioned above, one of the current challenges in the aneuploidy field is to generate experimental systems that recapitulate the chromosomal gains and losses found at the somatic or germline level in cancer or in congenital aneuploidy syndromes. As a proof of principle of this application, we used KaryoCreate to model frequent aneuploidy events found in colorectal cancer. Analysis of the TCGA dataset showed that chromosome 18q loss is observed in about 62% of colorectal cancer patients, being the second most frequent event, preceded by chromosome 20 gain (Fig. 5A). Importantly, based on the TCGA dataset we found that the presence of 18q loss is associated with worse survival in colorectal cancer patients (p=0.04, logrank test, Fig. 5B). Chromosome 7 gain is also a frequent event, present in about 50% of patients (Fig. 5A).

**Figure 5.**
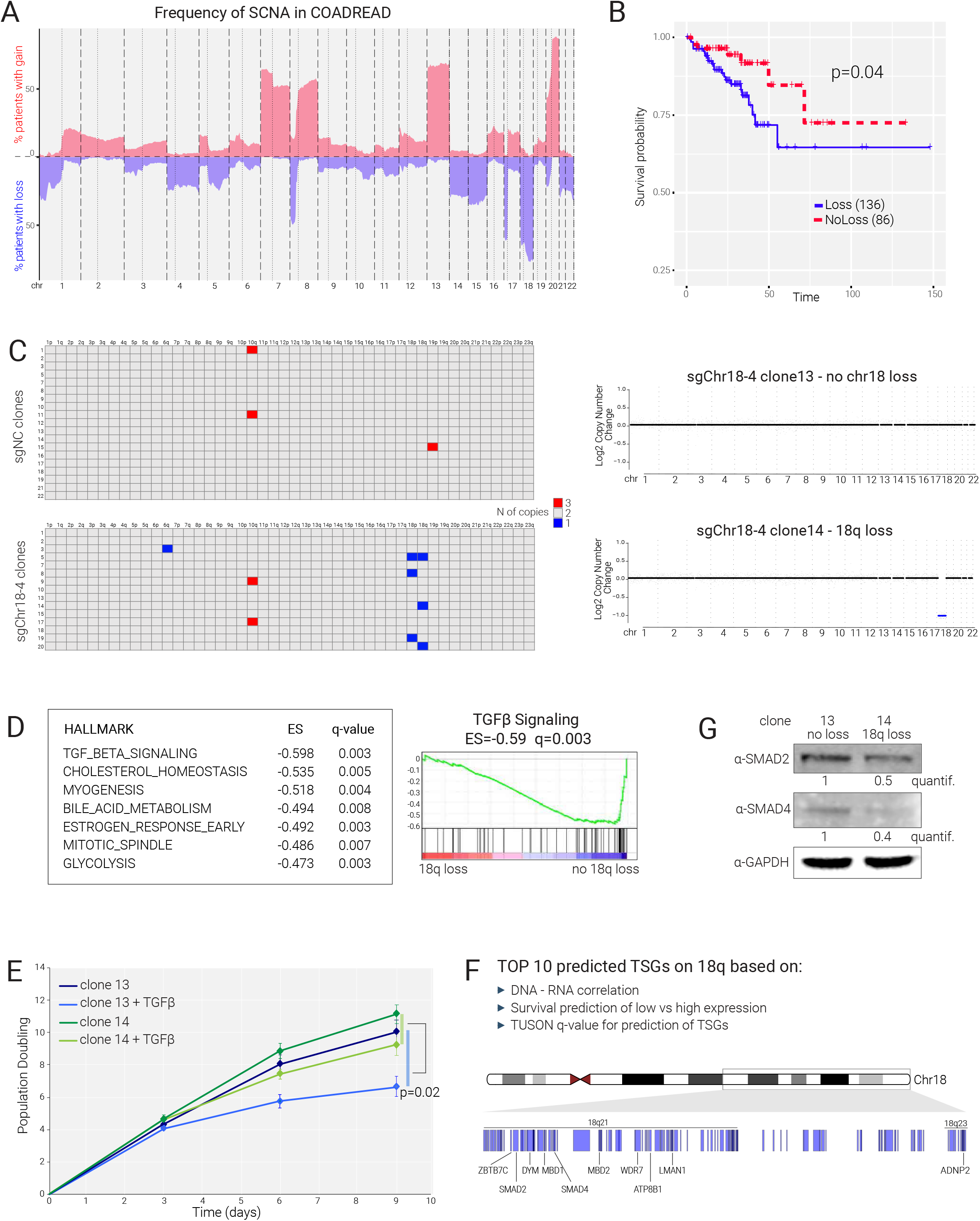
Loss of 18q in colon cells promotes resistance to TGFβ signaling. (A) Frequency of copy number alteration in colorectal cancer (TCGA – COADREAD). The percentage of patients with gain or loss for each chromosome is shown. (B) Survival analysis (Kaplan-Meier curve) for colorectal cancer patients ((TCGA – COADREAD) displaying or not 18q loss. (C) KaryoCreate using sgNC, or sgChr18-4 was performed on hCEC hTERT p53-/- diploid cells and single cell-derived clones were analyzed by WGS to identify arm-level gains and losses. Plots of copy number alterations of two representative clones from the cells treated with sgChr18-4 are shown on the right. (D) Cells were treated as in (C) and bulk RNAseq was performed on clones 13 and 14 (duplicates). Differential expression analysis was performed between clone 14 and clone 13 using DESeq2 and gene set enrichment analysis was performed using the Hallmark gene sets; the top 7 pathways enriched in clone 14 are shown including TGFβ signaling as the top enriched pathway. (E) Cells were treated as in (C) and the clones 13 and 14 were cultured in the presence of TGFβ (20ng/ml) for 9 days. Cells were counted every three days in duplicates. P-value is derived from the Wilcoxon test comparing the difference in cell number between clone 14 treated and clone 14 untreated versus the difference in cell number between clone 13 treated and clone 13 untreated. (F) Top 10 predicted tumor suppressor genes (TSG) on 18q and their location. TSG located on 18q were predicted based on three gene-based parameters (TCGA dataset): correlation between DNA and RNA level of the gene; survival analysis comparing patients with high versus low expression of the gene; TUSON-based q-value for the prediction of TSGs based only on point mutations. See text for details. (G) Cells were treated as in (C) and cWestern blot analysis for SMAD2 and SMAD4 levels in clones 13 and 14, with were analyzed by western blot for the amount of SMAD4, SMAD2 and GAPDH as control. Quantification is shown below SMAD2 and SMAD4.

To model these events, we performed KaryoCreate on hCEC TP53-/- cells using sgRNAs sgChr7-1, sgChr18-4, or sgNC as described above (see **Methods**). Single cell clones (20-23) were derived and evaluated for copy number profile by low-pass whole genome sequencing. Strikingly, compared to clones derived from the sgNC control condition, clones derived from sgChr7-1 showed an increase in the frequency of chr7 chromosome- or arm-level gains (from 0% in sgNC to 22% in sgChr7-1) but not of chr7 losses (0% in both conditions) (Fig. 5C), while clones derived from sgChr18-4 showed an increase in frequency of chr18 chromosome- or arm-level losses (from 0% in sgNC to 30% in sgChr18-4) but not of gains (0% in both conditions) (**Fig. S5B**). This finding recapitulates the recurrent patterns seen in human tumors, where chromosome 18 is frequently lost but virtually never (2%) gained while chromosome 7 is frequently gained and almost never (0.3%) lost. Importantly, we did not observe a change in the frequency of gains or losses of the chromosomes that were not targeted by the specific sgRNA. We noticed the presence of 10q gain in ∼20% of the clones derived in all three conditions (sgChr7-1, sgChr18-4 and sgNC), likely representing an event present in about 20% of the cells in the initial population.

Given its association with poor survival (Fig. 5B), we set out to characterize the phenotypic features of isogenic clones containing or not chromosome 18q loss. We started with two clones derived from the hCEC KaryoCreate sgChr18-4, one disomic for chromosome 18 (clone 13) and one with loss of one of the two copies of 18q (clone 14). We performed bulk RNA-seq analyses of these two clones in duplicates and performed differential expression analysis using DESeq2 (Love et al., 2014). Gene set enrichment analysis (GSEA) for cancer hallmarks showed that the top pathway downregulated in clone 14 compared to clone 13 was TGFβ signaling (Enrichment Score=-0.59; q-value=0.006), followed by cholesterol homeostasis, myogenesis and bile acid metabolism (Fig. 5D). TGFβ normally inhibits proliferation of colon epithelial cells by promoting their differentiation and its inhibition through intestine niche factors like Noggin is essential to allow proliferation and expansion of colon epithelial cells (Massagué et al., 2000). Since TGFβ signaling was the top pathway most significantly decreased in cells with 18q loss compared to control cells, we tested the effect of TGFβ activation in our clones. We performed an in vitro cell proliferation assay where we cultured clones 13 and 14 in the presence of TGFβ (20ng/ml) for 10 days. While TGFβ treatment reduced the growth of clone 13 cells by about 45% at day 9, TGFβ treatment reduced cell growth on clone 14 by less than 10% at the same time point (Fig. 5E; p=0.02). Altogether, these data suggest that 18q deletion leads to a decrease in the ability of the cells to respond to the growth inhibitory signal derived from TGFβ treatment. This result was also recapitulated for clone 5 that lost the entire chromosome 18, where no significant reduction in cell growth was found after TGFβ treatment (data not shown).

Chromosome 18q harbors an important tumor suppressor gene, SMAD4 (located on 18q21.2), which encodes for a transcription factor that is critical for mediating response to TGFβ signaling (Drost et al., 2015; van de Wetering et al., 2015). SMAD4 is commonly deleted in colorectal cancer as a consequence of chromosome 18q loss. In fact, SMAD4 deletion that is independent of chromosome 18q loss only occurs in 4% of patients. SMAD4 harbors point mutations in 29% of patients and thus 18q loss may serve to abolish SMAD4 function through loss of the remaining wild-type allele. However, it is unknown if the decreased survival in patients with 18q loss (Fig. 5B) is a consequence of the complete loss of SMAD4 (point mutation and deletion of other allele) or a consequence of simultaneous loss of several potential tumor suppressor genes residing on 18q, as others have suggested (Thiagalingam et al., 1996). To distinguish between these two possibilities, we assessed the contribution of 18q loss to the probability of patients’ survival after excluding patients with point mutations in SMAD4. If 18q loss serves to abolish SMAD4 function through deletion of the wild-type allele, we would predict that 18q loss would lose its association with patient survival after patients with SMAD4 mutations are excluded. Interestingly, 18q loss was still a significant predictor of survival after removing these patients, indicating that decreased patient survival could be a consequence of deletion of several tumor suppressor genes on 18q (**Fig. S5 C**, p-value of 0.006, lower than in the analysis that included all patients, see Fig. 5B).

To systematically predict tumor suppressor genes located on 18q, we developed a score for all the genes on 18q using three computational parameters based on the TCGA-COAD dataset: 1. The correlation between DNA and RNA level of each gene across patients, indicating whether a DNA copy number change was reflected at the RNA level; 2. The association of expression level of each gene with patients’ survival; 3. TUSON-based prediction of the likelihood for a gene to behave as a tumor suppressor gene based on its pattern of point mutations ((Davoli et al., 2013); see also **Methods**). The top ten genes predicted by this method were (in order of ranking position) SMAD2, ADNP2, MBD1, ATP8B1, WDR7, MBD2, DYM, SMAD4, ZBTB7C and LMAN1 (Fig. 5F). SMAD2, a paralogue of SMAD4, is found on 18q21.1 and is also a transcription factor that acts downstream of TGFβ signaling (Eppert et al., 1996; Massagué et al., 2000). Thus, the co-deletion and concomitant decrease in gene dosage of both SMAD4 and SMAD2 could synergistically mediate the unresponsiveness of cells to TGFβ signaling.

We set out to test the role of decreased dosage of SMAD2 and SMAD4 in our clone containing 18q loss. We confirmed by both RNAseq and by WB a decrease in both SMAD2 and SMAD4 in clone 14 compared to clone 13 (Fig. 5G; SMAD4 Log2FC: −0.78; p<0.0001; SMAD2 Log2FC: −0.75; p<0.0001). Furthermore, when we overexpressed SMAD2, SMAD4 in clone 14 cells, we found a significant decrease in proliferation rate after TGFβ treatment (**Fig. S5D**). Overexpression of both SMAD2 and SMAD4 had a synergistic effect leading to further decrease in proliferation, similar to clone 13 (**Fig. S5D**). Altogether, our computational and experimental data suggest that chromosome 18q loss, one of the most frequent events in gastro-intestinal cancers, is associated with poor survival and promotes resistance to TGFβ signaling likely due to the synergistic effect of simultaneous deletion of haploinsufficient genes.

## DISCUSSION

While much progress has been made to decipher causes of somatic and germline aneuploidy and consequences of aneuploidy in model systems (primarily yeast and mouse models), our knowledge of the consequences of specific chromosomal gains and losses in human cells is still limited. This is due in part to the lack of systems available to engineer isogenic genetic models of aneuploidy. In this work, we describe KaryoCreate, a novel method to induce chromosome-specific gains and losses in human cells by targeting chromosome-specific centromeric α-satellite arrays. We demonstrate induction of aneuploidy with an average frequency of 8% for gains and 12% for losses across 9 different human chromosomes. This method could be extended to all human chromosomes in principle, with the exception of chromosome Y due to the lack of sgRNAs with a sufficiently high number of binding sites. Using KaryoCreate, we successfully generated cell clones with chromosome-specific gains or losses that are stable for several population doublings. Furthermore, we showed that chromosome 18q loss, one of the most frequent aneuploidies in gastro-intestinal tumors, promotes resistance to TGFβ signaling.

### Chromosome-specific centromeric sgRNAs

Using the latest genome assembly from the T2T consortium, we were able to design sgRNAs binding uniquely to one specific centromere (Nurk et al., 2022). Since centromere sequences are variable across the human population, we designed sgRNAs using two genome assemblies (CHM13 and GRCh38) and tested them in one or two cell lines (hCEC, RPE), increasing the likelihood that the designed sgRNAs target conserved regions across the human population and thus can be applied to other cell lines. Among the 75 tested, we validated 24 sgRNAs targeting in a specific manner 15 different chromosomes by imaging (Fig. 1F). For several of the remaining chromosomes, we showed that the induction of gains and losses is still possible using one sgRNA targeting multiple chromosomes.

Our data also shed light on the design and use of sgRNAs for human centromeres. First, while sgRNA efficiency (by imaging) correlated with the number of binding sites, there was no correlation between the sgRNA efficiency and the activity score predicted using a commonly used algorithm for designing sgRNAs (Doench et al., 2016). This suggests that the set of rules that define efficient sgRNAs at the centromeres may be substantially different than the ones that apply to the rest of the genome; additional studies are needed to confirm this finding and identify those rules. Moreover, our data suggest that using more than one sgRNA does not improve the efficiency of aneuploidy induction (compared to single sgRNA) using KaryoCreate (**Fig. S3E, F**). Because of the repetitive nature of centromeric HOR, any pair of sgRNAs is predicted to bind multiple times and in relatively close proximity, potentially inducing competition or interference among KNL1^Mut^-dCas9 molecules. Finally, It has been previously shown that the size of centromeres can dictate the propensity of each chromosome to undergo missegregation (Dumont et al., 2020). In our case, the differential efficiency in chromosome targeting using KaryoCreate is likely due to different efficiency of the sgRNAs across chromosomes, although future studies are required to fully address this question in the context of our system.

### Comparison of KaryoCreate with other methods

The importance of generating cell systems to model aneuploidy is underlined by the increasing number of methods that have been recently developed to induce chromosome-specific missegregation, especially chromosome losses, as described above (see Introduction) (Rayner et al., 2019; Taylor et al., 2018; Zuo et al., 2017). Most of the methods to obtain chromosome losses involve the use of catalytically active Cas9 to eliminate chromosomes or parts of chromosomes, introducing DNA double strand breaks and potentially instigating undesired genomic instability (see for example (Rayner et al., 2019)). Of particular interest are two methods that were recently described to induce chromosome-specific gains and, mostly, losses (Tovini et al., 2022; Truong et al., 2022). Both methods target non-centromeric repeats and have been so far successful for two chromosomes: chromosome 1, using a sub-telomeric repeat, and chromosome 9 using a peri-centromeric repeat. Specifically, the work from Tovini et al. uses nuclease-dead Cas9 fused to the kinetochore-nucleating domain of CENPT to form an ectopic kinetochore. The data suggests activation of the spindle assembly checkpoint and requires a transient treatment with MPS1 inhibitor (such as revesine) to allow mitosis to progress upon missegregation. Truong et al. artificially tethered a plant kinesin to pull the chromatids towards one pole of the mitotic spindle. In this method, the introduction of a kinesin to a non-centromeric region generates the formation of a potential pseudo-dicentric chromosome pulled to opposite sides of the spindle, as suggested by the fact that most aneuploidies observed are losses of part of the targeted chromosome (chromosome 9). While both methods will be useful to dissect the spindle assembly checkpoint, the fate of dicentric chromosomes and the biophysical properties of chromosomal behavior at the metaphasic plate, KaryoCreate is profoundly different. We aimed to generate a method that would be applicable to potentially any chromosome by relying on endogenous centromeric sequences (present on all chromosomes) allowing the generation of (almost) any karyotype of interest. Also, we wanted to induce aneuploidy in the most physiological setting and avoid any use of mitotic drugs. Our data suggest that cells progress normally through the cell cycle with an expected delay in metaphase probably due to attempts in correcting merotelic attachments (Cimini et al., 2001; Gregan et al., 2011). Overall, we do not observe any defects in cell division or major indications of unexpected chromosomal instability both by metaphase spread analysis and sequencing (e.g. dicentric or focal amplifications), apart from arm events (see below). Notably, compared to existing technologies Karyocreate has a high capability of inducing specific aneuploidies. In the recent study by (Truong et al., 2022) the authors applied their method to chromosome 9 and showed the induction of arm-level losses in about ∼16% of cells for 9q and∼3% for 9p, and arm-level gains of∼3% for 9q and <1% for 9p; it was unclear whether any whole chromosome loss or gain was induced, likely due to the formation of dicentric chromosomes. Although it is very to hard to compare methods especially due to the use of different cell lines, the efficiency of induction of chromosome gains and losses for KaryoCreate was on average 9% for arm-level gains, 13% for arm-level losses, 8% for whole chromosome-chromosome gains and 11% for whole-chromosome losses. Furthermore, KaryoCreate showed efficiency in targeting multiple chromosomes at the same time, expanding its application to combinations of chromosomes.

### Targeting centromeric *α-satellites* to engineer chromosome-specific aneuploidy

We show that tethering of chimeric proteins comprising mutant forms of KNL1 and dCas9 to human centromeres induces chromosome and arm-level gains and losses. Of the two KNL1 mutant proteins that we tested, we initially predicted that KNL1^S24A;S60A^-dCas9 would result in a more efficient induction of chromosome gains and losses than KNL1^RVSF/AAAA^-dCas9, owing to a more efficient inhibition of Aurora B-mediated error correction through recruitment of PP1; however, this was not the case. There are many possible reasons, including protein stability: KNL1^RVSF/AAAA^-dCas9 levels tend to be generally higher than KNL1^S24A;S60A^-dCas9. NDC80-CH1-dCas9 and NDC80-CH2-dCas9 constructs showed very low protein expression (**Fig. S2C**), but relatively high induction of chromosome gains and losses (**Fig. S3B**); thus it is plausible that specific modifications to render those proteins more stable may render the system even more efficient. Future studies will be necessary to solve these questions and to further improve the efficiency of the technology.

Importantly, while combining multiple sgRNAs does not improve efficiency, boosting the expression of KNL1^Mut^-dCas9 through lentiviral-mediated delivery (e.g. inducible promoter) and FACS sorting based on cell surface markers present on the chromosome of interest allows enriching for the cell population containing the desired chromosome gain or loss (**Fig. S3B-E**). Our data suggest that engineering new cell lines for KaryoCreate may require testing of the different constructs to achieve the highest expression levels for greater results.

### Induction of arm-level gains and losses

About 55% of the aneuploidy events generated by KaryoCreate are arm-level events. In addition, we observed more losses (60%) compared to gains (40%) for both chromosome and arm events. Single cell-derived clones containing specific arm-level losses (e.g. chromosome 18q loss) can also be generated from the polyclonal population (Fig. 5C).

The induction of arm-level events may be due to several reasons. Some centromeres (for example the one on chromosome 7) can contain two juxtaposed HOR arrays that can both support centromere assembly, being structurally arranged as a dicentric chromosome (Maloney et al., 2012). It is possible that the recruitment of KNL1^Mut^-dCas9 to centromeres of structurally dicentric chromosomes, by preventing error correction, may lead to unresolved merotelic attachments that could result in DNA breaks at the centromere, generating arm-level aneuploidy. Nevertheless, the presence of multiple active HOR is not predicted to be a widespread feature across chromosomes (Maloney et al., 2012), while we observe the induction of arm-level events for all chromosomes that we have tested. Another possibility is the fact that the mere recruitment of a bulky protein to the centromere may influence transcription and/or replication thus causing breaks and centromeric instability (Barra and Fachinetti, 2018; Bury et al., 2020; Sullivan and Sullivan, 2020; Whinn et al., 2019). Additional studies are needed to better understand how dCas9 affects chromosome segregation when recruited to the highly repetitive centromeric regions. Overall, the possibility of KaryCreate to induce arm events in addition to whole chromosome changes is an additional advantage of the technology, as several tumors present this type of aneuploidy.

### Chromosome-specific aneuploidy as drivers of cancer hallmarks

In this study we applied KaryoCreate as a robust tool to model and study cancer aneuploidy. When we used KaryoCreate on colon epithelial cells (hCEC) to generate single cell derived clones, we showed that chromosomes that are frequently gained (chromosome 7) or lost (chromosome 18) in colorectal tumors, tend to also be gained or lost in the generated clones (Fig. 5C, **Fig. S5C**). This suggests that the selective pressure acting during tumor evolution to shape recurrent patterns of aneuploidy may be also acting *in vitro* to promote survival of cells with specific gains or losses (Davoli et al., 2013; Watkins et al., 2020). Furthermore, we validated that aneuploidies generated by KaryoCreate are stable over time, as shown for the hCEC clones with 18q loss. We showed that chr18q loss can promote resistance to TGFβ signaling in colon cells, a cancer-associated phenotype. While SMAD4 is an important tumor suppressor gene located on 18q, our data indicate that the phenotype determined by 18q loss may be due not only to one gene loss but to the cumulative effect of losing multiple tumor suppressor genes. Using a series of parameters based on the gene mutation pattern, specifically whether the RNA level changes proportionally to the DNA copy number and whether the RNA-based gene level predicts patients’ survival, we predicted several potential tumor suppressor genes on 18q in colorectal cancer. Experimentally, we tested SMAD4 and SMAD2 for their effect on TGFβ treatment; our data suggest that a 50% decrease in gene dosage of these genes leads to a significant and synergistic effect on the TGFβ signaling pathway.

Previous studies have proposed that a single cancer driver gene may determine a strong impact on the phenotypic effect of a chromosome gain or loss (McFadden et al., 2014; Trakala et al., 2021). Other studies, including previous work on chromosome 18, have proposed that it is not a single gene that mediates the selective advantage imparted by a specific aneuploidy but it is the cumulative effect of multiple genes (Thiagalingam et al., 1996;Davoli et al., 2013; Thiagalingam et al., 1996; William et al., 2021; Xue et al., 2012). In line with previous human genetics analyses (Thiagalingam et al., 1996), we predicted multiple haplo-insufficient tumor suppressor genes localized on 18q that are likely to act cumulatively in tumor progression. Furthermore, if SMAD4 was the only critical tumor suppressor localized on 18q and the main function of 18q was to delete the remaining WT copy, leading to a homozygous loss of SMAD4, we predict that 18q loss would predict survival in colorectal cancer patients only when considering patients with SMAD4 mutations. Notably, this is not the case: after we exclude the patients with SMAD4 point mutations, 18q loss was still able to predict patient survival (**Fig. S5C**). In addition, we found that reduction of SMAD4 by itself was not responsible for the increase in resistance to TGFβ signaling, but it also required deletion of SMAD2, also located on 18q. Thus, chromosome 18 loss may drive this phenotype by hemizygous deletion of two haploinsufficient genes acting in the same pathway. Furthemore, we speculate that 18 loss may drive other cancer-related pathways (that appeared to be altered in our gene expression analysis) through the deletion of additional haploinsufficient tumor suppressors on 18q potentially acting in similar pathways.

### Future directions

Altogether, these data describe KaryoCreate as a novel powerful resource for the scientific community to foster our understanding of chromosome missegregation and aneuploidy in different fields of biomedicine, including genetics, centromere biology, cancer aneuploidy and congenital aneuploidy (Sheppard et al., 2012). Chromosome-specific sgRNAs will allow studying the biology of human centromeres at single-chromosome resolution. For example, our system will allow live-cell imaging of individual centromeres or groups of centromeres in human cells, using fluorescently-tagged dCas9. Furthermore, the direct study of abnormal gene dosage in human congenital syndromes and cancer will improve the identification of critical genes that are drivers of specific phenotypes and may foster the development of specific therapies.

## ACKNOWLEDGEMENTS

NYU Langone’s Genome Technology Center is supported by the Cancer Center Support Grant P30CA016087 at the Laura and Isaac Perlmutter Cancer Center. We acknowledge all the members of the Davoli lab and Jef Boeke for helpful insights and discussion. We thank Liam Holt and Gregory Brittingham for help with the live-cell imaging experiments. We thank Jennifer DeLuca for generously providing the NDC80 aa1-207 construct and Karen Miga for the initial computational validation of a few sgRNAs. The computational requirements for this work were supported in part by the NYU Langone High Performance Computing (HPC) Core’s resources and personnel. Teresa Davoli and members of her lab are supported by the Cancer Research UK Grand Challenge and the Mark Foundation for Cancer Research (C5470/A27144), R37 R37CA248631 and the MRA Young Investigator Award. Sarah Keegan and David Fenyo were supported by grant U24CA210972 from the National Cancer Institute’s Clinical Proteomic Tumor Analysis Consortium, and contract S21-167 from Leidos Biomedical Research, respectively. Lizabeth Katsnelson, Aleah Goldberg and Joseph Mays were supported by a Cell Biology training grant T32 GM136542.

## METHODS

### RESOURCE AVAILABILITY

#### Lead contact

Further information and requests for reagents may be directed to the lead contact, Teresa Davoli.

#### Computational sgRNA prediction

We downloaded the CHM13 centromere and whole genome reference from T2T Consortium (https://sites.google.com/ucsc.edu/t2tworkinggroup/chm13-cell-line) (Nurk et al., 2022) and hg38 reference from UCSC genome browser. We selected the HOR region with the classification for “live” or “hor_L”. Then we split the HOR Live regions into different chromosomes. For each HOR Live region file, we searched all possible sgRNA binding sites with the specific pattern (20 nucleotides as sgRNA plus 3 nucleotides NGG as PAM). For each possible sgRNAs, we counted the frequency of each sgRNA binding sites for every centromere and the whole genome. We also calculated the GC content for each sgRNA.

In order to ensure chromosome-specificity of the sgRNAs, for each sgRNA we determined the chromosome specificity score, defined as ratio between the number of binding sites on the centromere of the target chromosome and the total number of sites across all centromeres (given as a fraction or as a percentage after multiplication by 100) as well as the centromere specificity score, defined as the ratio between the number of binding sites on the centromere of the target chromosome and the number of binding sites across the whole genome (given as a fraction or as a percentage after multiplication by 100).

The sgRNA efficiency (activity score) was evaluated using Doench’s method (Doench et al., 2014). In brief, a support vector machine (SVM) method was used to generate sets of features and a logistic regression was trained to discriminate the top sgRNA for each gene. The model was trained based on the data of the top genes and predicted the data for the remaining genes. The feature selection was generated based on the nested stratified cross validation. After validation, we trained a final model using the data from different resources, which used 72 features from Doench’s dataset (Doench et al., 2014) and 9 features from our previous dataset.

The feature presented in both dataset can be used to calculate the efficiency of sgRNA (activity score). In other words, if we let the model weights for the features *i* for a particular guide *s_j_* be *w_ij_*, let the intercept be int. Then the activity Score *f*(*s_j_*) is given via logistic regression as:

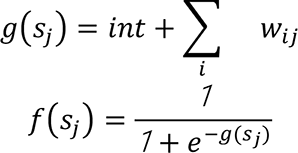

Activity score *f*(*s*_j_) would fall into the range [0,1], the worst sgRNA for a chromosome would receive a score of 0 and the best sgRNA would have a score approaching 1. For the chromosome Y, since the CHM13 is a female cell line, all the binding sites are evaluated based on hg38.

### EXPERIMENTAL METHODS

#### Cell Lines

All cells were grown at 37°C with 5% CO2 levels. hCEC (human colonic epithelial cells) (Ly et al., 2011) were cultured in a 4:1 mix of DMEM:Medium 199, supplemented with 2% FBS, 5 ng/mL EGF, 1 μg/mL hydrocortisone, 10 μg/mL insulin, 2 μg/mL transferrin, 5 nM sodium selenite, pen-strep, and L-glutamine. RPE-1 (human retinal pigment epithelial-1) cells (Maciejowski et al., 2015) and HCT116 (human colorectal carcinoma-116) cells were incubated in DMEM, supplemented with 10% FBS, pen-strep, and L-glutamine. For long-term storage, cells were cryopreserved at −80°C in 70% medium (according to cell line), 20% FBS, 10% DMSO. TP53 was knocked-out in hCEC by transfection with a Cas9 containing plasmid (Addgene #42230) and plentiGuide-Puro expressing the following sgRNA: GCATGGGCGGCATGAACCGG. Clones were derived and tested for the expression of TP53.

#### Cloning of KaryoCreate Constructs and 3xmScarlet-dCas9

Cas9 and dCas9 without ATG and without stop codon (for N-terminal and C-terminal tagging respectively) were cloned into D-TOPO vector (Thermo #K240020). Cloning of KNL1^RVSF/AAAA^-dCas9 was achieved by inserting KNL1 PCR product (aa1-86, amplified from Addgene plasmid #45225, (Liu et al., 2010)) into XhoI-digested pENTR-dCas9 (no ATG) using Gibson assembly. The GGSGGGS linker was added between KNL1 and dCas9 by including the coding nucleotide sequence in the reverse primer for the KNL1 PCR product. Cloning of KNL1^S24A;S60A^-dCas9 was achieved starting from KNL1^RVSF/AAAA^-dCas9 and inserting the appropriate mutations using Gibson assembly.

Cloning of NDC80-CH1-dCas9 was achieved by Gibson assembly of NDC80 aa1-207 (generously provided by Dr. Jennifer DeLuca) with BamHI-digested pENTR dCas9 (ATG). Cloning of NDC80-CH2-dCas9 was achieved in a similar way except that 2 CH domains were cloned in tandem separated by a linker.

All pENTR vectors were cloned into specific pDEST vectors by LR reaction (Thermo #11791020) following the manufacturer’s instructions. pDEST vectors used in this study are: pHAGE (Blast resistance) or pINDUCER20 (or pIND20, neomycin resistance) (Meerbrey et al., 2011).

To generate an additional inducible KNL1^Mut^-dCas9 expression vector (in addition to pIND20) the FKBP12 degradation domain (DD, Banaszynski 2006) was first amplified from Degron-KI-donor backbone (Addgene #65483) and inserted at the N-terminus of the fusion protein sequence in pENTR-KNL1-dCas9 using Gibson cloning. Gateway LR cloning was then used to yield the expression vector, pHAGE-DD-KNL1^Mut^-dCas9.

pHAGE-3xmScarlet-dCas9 plasmid was generated by three-step cloning. (1) Single mScarlet with different flanking overhangs were made by PCR amplification of the plasmid containing mScarlet sequence. Three mScarlets were assembled in series and inserted into the pAV10 vector by Golden Gate cloning technology (BsaI enzyme). (2) 3xmScarlet with homology regions at two ends for Gibson reaction in the next step was generated by PCR amplification of pAV10-3xmScarlet plasmid created in the previous step (forward primer: AGGCTCCGCGGCCGCCCCCTTC, reverse primer: CCAGGCCGATGCTGTACTTCTTGTCCGACC). DNA fragment of the correct size was purified from agarose gels. (3) 3xmScarlet was assembled with linearized pENTR-dCas9 (XhoI enzyme) by Gibson reaction to form pENTR-3xmScarlet-dCas9 plasmid. Then 3xmScarlet-dCas9 was transferred into pHAGE destination vector by Gateway cloning method.

#### Cloning of sgRNAs

We modified plentiGuide-Puro (Addgene #52963) by Gibson assembly to contain a modified scaffold sequence to improve efficiency, obtaining pLentiGuide-Puro-FE (Sanjana et al., 2014). sgRNAs were designed and cloned into this pLentiGuide-Puro-FE vector according to the Zhang Lab General Cloning Protocol ((Ran et al., 2013), also https://www.addgene.org/crispr/zhang/). To be suitable for cloning into *BbsI*-digested vectors, sense oligos were designed with a CACC 5’ overhang and antisense oligos were designed with an AAAC 5’ overhang. The sense and antisense oligos were annealed, phosphorylated and ligated into either *BbsI*-digested pLentiGuide-Puro-FE for KaryoCreate and imaging purposes or pX330-U6-Chimeric_BB-CBh-hSpCas9 (Addgene #42230) for CRISPR/Cas9 editing applications. Sequences were confirmed by Sanger sequencing.

#### Lentivirus production and nucleofection of hCEC cells

For transduction of cells, lentivirus was generated as follows: one million 293T cells were seeded in a 6-well plate 24 hours before transfection. The cells were transfected with a mixture of gene transfer plasmid (2 μg) and packaging plasmids including 0.6 μg ENV (VSV-G), 1 μg Packaging (pMDLg/pRRE), and 0.5 μg pRSV-REV along with CaCl2 and 2xHBS or using lipofectamine (Thermo #L3000-075).The media was changed 6 hours later and virus was collected 48 hours after transfection by filtering the media through a 0.45 μm filter. Polybrene (1:1000) was added to filtered media before infection.

Nucleofection of hCEC cells was carried out using the Amaxa Nucleofector II (Lonza), optimized for the HCT116 cell line. Approximately 1 million cells suspended in 100 μL of electroporation buffer (80% 125 mM Na2HP04.7H20, 12.5 nM KCl, 20 % 55 mM MgCl2) were subjected to electroporation in the presence of a vector, and immediately returned to normal medium.

#### Western blot analysis

Cells were harvested by trypsinization, lysed in 2X NuPAGE LDS buffer (Thermo #NP0007) at 10^6^ cells in 100μl of buffer. DNA was sheared using a 28 1/2-gauge insulin syringe and lysate was denatured by heating at 80°C for 10 minutes. Lysate equivalent to 10^5^ cells was resolved by SDS/PAGE using NuPAGE 4-12% Bis-Tris mini Gel and transferred to a PVDF membrane. The membrane was then blocked in 5% milk in TBS with 0.1% Tween-20 (TBS-T) for 1 hour at room temperature. Afterward, the membrane was probed with Cas9 (Abcam #ab191468)(1:1000) and GAPDH (Santa Cruz #sc-47724)(1:10000 or 1:100000) primary antibodies and incubated in 2.5% milk in TBS at 4°C overnight.

Subsequently, the membrane was washed three times with TBS-T and incubated with HRP-anti-Mouse secondary Ab (Abcam #ab205719)(1:1000) in 2.5% milk/TBS for 1 hour at room temperature. Signals were detected using an ECL system using 1:1 detection solution after three 10-minute washes in TBST. Images were acquired using a BIORAD transilluminator.

#### Fluorescence in situ hybridization (FISH)

FISH analysis was carried out on interphase nuclei and metaphase spreads. Cells at 70% confluence were harvested by trypsinization (after 3-4 hours treatment with 100 ng/mL colcemid [Roche #10295892001] for metaphase spreads), washed with PBS, suspended in 0.075 M KCl at 37°C, and fixed in methanol-acetic acid (3:1) at 4°C. Fixed cells were dropped onto glass slides and they were allowed to air dry overnight. The slides were next incubated with RNase solution (20 μg RNase A in 2xSSC) for one hour at 37°C in a dark moist chamber. Denaturing was performed using a 70% Formamide solution (in 2xSSC) for three minutes at 80°C prior to hybridization. Biotinylated/Digoxigeninated probes were obtained by nick translation from BAC DNA (RP11-22N19 for chromosome 7, RP11-76N11 for chromosome 13, and RP11-787K12 for chromosome 18 from the BACPAC Resource Center). 200ng of each labeled probe, together with 8 μg Human Cot-I DNA (Thermo #15279011) and 3 μg Herring Sperm DNA (Thermo #15634017) were precipitated for one hour at −20°C in 1/10 volume of 3M Sodium Acetate and 3 volumes Ethanol. The pelleted probe was washed with 70% ethanol, air dried, and resuspended in hybridization solution (50% Deionized Formamide, 10x Dextran Sulfate =, 2=xSSC). The hybridization solution containing the probes was then denatured at 80°C for 10 minutes and then incubated at 37°C for 20 minutes to allow annealing of the Cot-I competitor DNA. The sealed hybridized slides were then incubated at 37°C in a dark moist chamber overnight. The following day, slides were washed in 1xSSC at 60°C (3 times, 5 minutes each) and incubated with blocking solution (BSA, 2xSSC, 0.1% Tween-20) for 1 hour at 37°C in a moist chamber. Following blocking, the slides were incubated with detection solution containing BSA l, 2xSSC, 0.1% Tween-20, and FITC-Avidin conjugated 0.5 μl (Thermo #21221), and Rhodamine-AntiDigoxigenin 10 μl (SIGMA #11207750910) to detect the biotin and digoxigenin signals. Finally, slides were washed 3 times (5 mins each) with 4xSSC and 0.1% Tween-20 solution at 42° C. Slides were finally mounted with DAPI for DNA stain (Vector Laboratories #H-1200-10).

Images were acquired by a fluorescent microscope, Invitrogen^TM^ Evos^TM^ M700 imaging system or Nikon TI Eclipse. The number of fluorescent signals was counted in 100 intact nuclei per slide. Adobe Photoshop was used to count the signals and correct the images.

#### Live-cell imaging

Cells were plated on 35mm glass-bottom microwell dishes (MatTek P35G-1.5-14-C) one day prior to imaging. Imaging was performed at 37°C and 5% CO2 using an Andor Yokogawa CSU-X confocal spinning disc on a Nikon TI Eclipse microscope. Samples were exposed to 488 nm (30 ms) and 561 nm (100 ms) lasers and fluorescence was recorded with a sCMOS Prime95B camera (Photometrics). A 100X objective was used to acquire images at 0.9 µm steps (total range size=9 µm) every 1 or 3 minutes as indicated in the figure legends. Image analysis was performed using ImageJ and formatting (cropping, contrast adjustment, labeling) was performed in Adobe Photoshop.

#### Low-Pass Whole Genome Sequencing

Genomic DNA was extracted from trypsinized cells using 0.3 μg/μL Proteinase K (QIAGEN #19131) in 10mM Tris pH 8.0 for 1 hour at 55°C, then heat inactivated at 70°C for 10 minutes. DNA was digested using NEBNext® dsDNA Fragmentase® (NEB #M0348S) for 25 minutes at 37°C followed by magnetic DNA bead cleanup with Sera-Mag Select Beads (Cytiva #29343045), 2:1 bead to lysate ratio by volume. We created DNA libraries with an average library size of 320 bp using the NEBNext® Ultra™ II DNA Library Prep Kit for Illumina® (NEB #E7103) according to the manufacturer’s instructions. Quantification was performed using a Qubit 2.0 fluorometer (Invitrogen #Q32866) and the Qubit dsDNA HS kit (#Q32854). Libraries were sequenced on an Illumina NextSeq 500 at a target depth of 4 million reads in either paired-end mode (2 x 36 cycles) or single-end mode (1 x 75 cycles).

#### RNA bulk sequencing

Clones were plated in 6-well plates one day before collection. On the day of collection, cells were checked for confluency within 70-90% and normal morphology. Cells were washed twice with PBS and stored at −80C immediately. RNA was purified for bulk sequencing using the Qiagen RNeasy Mini Kit (Qiagen 74106). RNA concentration and integrity were assessed using a 2100 BioAnalyzer (Agilent, Santa Clara, CA). Sequencing libraries were constructed using the TruSeq Stranded Total RNA Library Prep Gold mRNA (Illumina, San Diego, CA) with an input of 250ng and 13 cycles final amplification. Final libraries were quantified using High Sensitivity D1000 ScreenTape on a 2200 TapeStation (Agilent, Santa Clara, CA) and Qubit 1x dsDNA HS Assay Kit (Invitrogen, Waltham, MA). Samples were pooled equimolar with sequencing performed on an Illumina NovaSeq6000 SP 100 Cycle Flow Cell v1.5 as Paired-end 50 reads.

#### Clone derivation

hCEC were transduced with pHAGE-DD-KNL1^Mut^-dCas9 and a sgRNA vectors and DD-KNL1^Mut^-dCas9 was stabilized with 100 nM Shield-1 (Takara, #632189) for 9 days. Three days after Shield-1 treatment, 20-500 cells were plated per 15 cm plate and were incubated in normal culture conditions until colonies were visible (∼2-3 weeks). Colonies were then picked by applying wax cylinders to the area surrounding each clone, trypsinizing the cells, and moving them to separate wells in 48-well plates for further expansion.

#### Single-cell RNA sequencing

scRNA-seq libraries were prepared using the 10X Chromium Single-Cell 3’ v3 Gene Expression kit according to the manufacturer’s instructions, including the manufacturer’s protocol for cell surface protein (hashtag antibody) feature barcoding. Up to 10 TotalSeq-B hashtag antibodies (BioLegend, San Diego, CA) were used for multiplexing samples in each sequencing run.

### QUANTIFICATION AND STATISTICAL ANALYSIS

#### Automated FISH counting

In addition to manual counting of FISH foci (shown in Figs 3 and S3), we also performed an automated FISH foci counting. In order to calculate the FISH counts automatically, we used an in-house developed python script, available publicly at https://github.com/davolilab/FISH-counting. Individual nuclei were segmented by applying an automatic threshold to the DAPI channel after a smoothing and contrast enhancement. Thresholded objects were filtered for area and solidity to remove erroneously segmented regions. For probe detection within segmented nuclei, a white tophat filter was applied to remove small spurious regions and then the “blob_log” function from scikit-image package (van der Walt et al., 2014) was utilized to identify and count fluorescent spots. Since it was observed that some FISH probes were incorrectly doubly counted, a distance cutoff was applied so that spots within a set (minimal) distance count as one. Then, we aggregated the probes numbers and calculated percentages for different spot counts. The script was run under Python 3.7 environment; for more details see the github repository.

#### Quantification of foci intensity

The regions corresponding to the FISH foci were determined by the threshold function of Fiji. Then the average intensity of each determined region was calculated as the representative of the brightness of the focus by Fiji (used in Fig S1E).

#### Low-pass whole genome sequencing analysis

Low-pass (∼0.1-0.5X) whole-genome sequencing reads of cells were aligned to reference human genome hg38 by using BWA-mem (v0.7.17) (Li and Durbin, 2009), followed by duplicate removal using GATK (Genome Analysis Toolkit, v4.1.7.0) (https://gatk.broadinstitute.org/hc/en-us) with default parameters to generate analysis-ready BAM files. BAM files were processed by the R Package CopywriteR (v1.18.0) (Kuilman et al., 2015) to call the arm-level copy numbers.

#### Bulk RNAseq analysis pipeline

RNA sequencing reads were processed, quality controlled, aligned, and quantified using the Seq-N-Slide software (Dolgalev, Igor, 2022). In short, total RNA sequencing reads were trimmed using Trimmomatic (Bolger et al., 2014) and mapped to the GENCODE human genome hg38 by STAR (Dobin et al., 2013). featureCounts (Liao et al., 2014) was used to quantify reads and generate a genes-sample counts matrix. Differential gene expression (DGE) analysis was completed with DESeq2 in R (Love et al., 2014). Gene ranks from DGE were used for pathway analysis using GSEA pre-ranked (Subramanian et al., 2005). Further plotting and statistical analyses were completed in R.

#### Single cell RNA Sequencing Data Pre-Processing

The CellRanger v6.1 pipeline (10X Genomics) was used to process single-cell RNA sequencing data. CellRanger count was used to align sequences and generate gene expression matrices. Sequences were aligned to a custom reference genome (designated 20210929-custom-ref) created with CellRanger mkref with default parameters. This custom reference was created using the pre-built GRCh38-2020-A human reference for CellRanger based on hg38, combined with sequences for dCas9, puromycin resistance, blastomycin resistance, and GFP {sources for sequences needed here}. Gene expression matrices were generated with each column representing a cell barcode and each row representing a gene or hashtag oligo sequences (HTO).

To identify the sample-of-origin for each cell barcode, the HTO count data from each 10X Chromium experiment were demultiplexed using the Seurat v4.0.3 package for R v4.1. Cell barcodes that could be confidently assigned to a single sample were kept. Several quality control thresholds were applied uniquely to each dataset on total gene number, total UMI counts, and total HTO counts to remove low-quality cells and potential cell doublets. Cells were also discarded if their proportion of total gene counts that could be attributed to mitochondrial genes exceeded 10%.

#### Modified CopyKat Analysis

A modified version of the CopyKat v1.0.5 pipeline for R was used to generate a Copy Number Alteration (SCNA) score for each chromosome arm in each cell. Samples from the same cell line in each 10X Chromium dataset were grouped together for analysis. Each such group of samples contained a diploid control sample used to set the SCNA value baseline centered around 0. For each analysis, genes expressed in less than 5% of the cells, HLA genes, and cell-cycle genes were excluded. The log-Freeman-Tukey transformation was used to stabilize variance and dlmSmooth() was used to smooth outliers. The diploid control sample for each set was used to calculate a baseline expression level for each gene. This value was subtracted from the samples in the set, centering the control sample expression around 0. Genes expressed in less than 10% of cells were then excluded from further analysis. The original CopyKat pipeline splits the transcriptome into artificial segments based on similar expression, and calculates a SCNA value for each segment. Instead, we generated a SCNA value for each chromosome arm by calculating the mean gene expression for the genes on that arm. A single SCNA value for the entire chromosome 18 was calculated instead for each arm of chromosome 18, due to its relatively small size. Gains or losses of a chromosome arm relative to the control sample (diploid) were called based on a threshold calculated from the control sample for each chromosome arm. The threshold is calculated as the

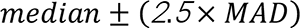

where the median is calculated from the SCNA values for each arm in the control sample, and the MAD (median absolute deviation) is calculated by the mad() function from the stats R package. Gains (or losses) are then called for a chromosome arm if its SCNA value is above (or below) the threshold for its sample set.

#### CopyKat Data Visualization

Heatmaps were generated using the ComplexHeatmap v2.8 R package {@Gu_2016_ComplexHeatmapR}. Each row represents one cell, each column represents a chromosome arm, and each value is the corresponding SCNA score. Column widths were scaled to the number of genes on the arm. For the heatmaps, cells were clustered by row of the chromosome of interest. Bar graphs were generated using the ggplot2 v3.3.5 R package.

#### Survival analysis

For survival analysis, the disease-free interval (DFI) and related clinical data was downloaded from cBioPortal (Liu et al., 2018). Arm-level copy number was downloaded from TCGA Firehose Legacy (https://gdac.broadinstitute.org). For each patient, purity *α*, ploidy *τ*, integer copy number *q*(*x*) *data* was downloaded from GDC (https://gdc.cancer.gov/about-data/publications/pancanatlas). Before the analysis, all the arm-level copy number *R*(*x*) were adjusted by the formula below:

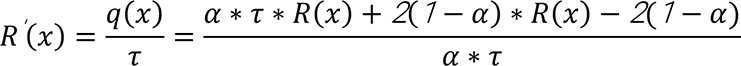

Patients with arm-level log2 ratio less than −0.3 would be regarded as an arm-level loss event. A log-rank test between the stratified patients and the Kaplan-Meier method was used to calculate the p-value and plot survival curves. Patients without clinical survival information were exclude from the analysis. In addition, a Cox proportional hazards (PH) regression model was also used to calculate each gene’s Hazard Ratio (HR) between the top 50% and bottom 50% expression.

#### Gene rank score analysis

For each gene on chromosome 18, we calculated the DNA-RNA Spearman’s correlation (rho value) from the TCGA-COAD READ dataset. Genes with no or very low frequency of SCNA (−0.02< DNA log2FC <0.02 in more than 70% of the patients) were removed because for those genes very little or no variance at the DNA level is likely to influence the correlations analyses.The association of expression level of each gene with patients’ survival was generated based on the HR from survival analysis.TUSON-based prediction algorithm of the likelihood for a gene to behave as tumor suppressor gene (TSG) based on its pattern of point mutation was from (Davoli et al., 2013) and was applied to the latest available TCGA dataset. A gene rank score was generated based on the sum rank of DNA-RNA correlation, hazard ratio from Cox proportional hazards regression and q-value from TUSON’s TSG prediction. A higher rank score means the gene is more important in this analysis.

## SUPPLEMENTARY MATERIAL

### LEGENDS TO SUPPLEMENTARY FIGURES

**Figure S1 (related to Fig. 1)**

A. Left Panel: Toxicity assay to test the efficiency of the centromeric sgRNAs. hCECs hTERT p53-/- expressing, or not, Cas9 were transduced with lentiviral vectors expressing the indicated sgRNAs. The percentage of live cells relative to EV was determined 7 days after transduction. Mean and standard deviation are shown; p-values are from Wilcoxon test (*=p<0.05). The sgRNAs that were or were not validated by imaging are shown. Right Panel: Western blot showing Cas9 expression.
B. Imaging efficiency for hCEC hTERT p53-/- cells (47, +7) expressing 3xmScarlet-dCas9 and sgChr7-1 as a polyclonal population and a derived clone (hCEC-clone#8). As compared to the polyclonal population, a higher percentage of cells showing the expected foci is present in clone 8. Western blot analysis for the expression level of 3xmScarlet-dCas9 in the polyclonal population and clone 8 is also shown on the right. Average frequency of cells displaying foci is shown for the polyclonal and clonal populations (<100 cells counted; in triplicates).
C. hCEC hTERT p53-/- cells transduced with a vector expressing a fluorescently-tagged dCas9 (3xmScarlet-dCas9 fusion) and with a vector expressing the indicated sgRNAs. Representative images of interphase cells are shown. See also Fig. 1F.
D. RPE cells were transduced with a vector expressing a fluorescently-tagged dCas9 (3xmScarlet-dCas9 fusion) and with a vector expressing the indicated sgRNAs. Representative images of interphase cells are shown.
E. Correlation between the intensity of the signal of the foci (measured with ImageJ/Fiji) and the number of predicted binding sites on the sgRNA efficiency score (Doench paper) (left panel) or the specific centromere based on the CH13 reference genome (right panel). Pearson correlation coefficients and corresponding p-values are shown.

**Figure S2 (related to Fig. 2)**

A. Map of different KaryoCreate constructs: dCas9, KNL1^Mut^; KNL1^S24A;S60A^ ; NDC80-CH1-dCas9 and NDC80-CH2-dCas9. The predicted function of each construct is also indicated on the right. See text for details.
B. Western blot showing the expression of the indicated construct in hCEC cells.
C. Western blot showing the expression of the indicated construct in hCEC cells where different mutant portions of KNL1 or NDC80 are fused at the N- or C-terminus of dCas9 (see also Fig. S2A). L:linker
D. hCEC hTERT p53-/- cells transduced with a vector expressing a fluorescently-tagged dCas9 (3xmScarlet-KNL1^Mut^-dCas9 fusion) and with a vector expressing sgChr7-1 or sgChr18-4.
E. Proliferation rate of hCEC hTERT p53-/- cells transduced with KNL1^Mut^-dCas9 and with the indicated sgRNAs.

**Figure S3 (related to Fig. 3)**

A. KaryoCreate experiment in hCEC hTERT p53-/- cells to compare the efficiency of different methods used to deliver KNL1^Mut^-dCas9. All the cells were transduced with sgChr7-1 and FISH quantification for chromosome 7 is shown. Methods: (1) pHAGE-KNL1^Mut^-dCas9 where this vector allowing constitutive expression is transfected in the cells; (2) pIND20-KNL1^Mut^-dCas9 where this vector is integrated in the cells and expression of the KNL1^Mut^-dCas9 is driven by Doxycycline treatment (1 μg/ml); (3) pHAGE-DD-KNL1^Mut^-dCas9 where expression of KNL1^Mut^-dCas9 is driven by treatment with shield-1 to stabilize the protein.
B. KaryoCreate experiment to compare the efficiency of different constructs in inducing chromosome gains and losses. hCEC hTERT p53-/- cells were transduced with sgChr7-1 and with all the indicated constructs. FISH quantification is shown for chr7 and chr18. Mean and standard deviation are shown; p-values are from Wilcoxon test (*=p<0.05).
C. hCEC hTERT p53-/- cells were transduced with KNL1^Mut^-dCas9 using different amount of virus (about 3 times more virus in the HIGH versus LOW sample, i.e. MOI of 6 for HIGH and of 2 for LOW). Western blot analysis for dCas9 is shown as well as the corresponding quantification (ImageJ).
D. hCEC hTERT p53-/- cells were treated as in C and then transduced with sgChr7-1. After 10 days, cells were processed for FISH using probes specific for chr7. Quantification of the % of cells with chr7 gains or losses for the indicated conditions is shown. Mean and standard deviation are shown; p-values are from Wilcoxon test (*=p<0.05).
E. hCEC hTERT p53-/- cells were transduced with KNL1^Mut^-dCas9 and with sgChr7-1 and/or sgChr7-3. Quantification shows the % of chr 7 gains or losses in each of indicated conditions.
F. hCEC hTERT p53-/- cells were transduced with KNL1^Mut^-dCas9 and with sgNC, sgChr9-3 and/or sgChr9-5. Quantification (by single cell sequencing) shows the % of chr 9 gains or losses in each of indicated conditions.
G. hCEC hTERT p53-/- cells were treated as in Fig. S3C-D using an MOI of 2 and FACS-sorted for the expression level of the cell surface protein EPHB4 whose gene is located on chr 7. FISH quantification shows the % of chr 7 gains or losses in each of indicated conditions. * refers to p-value<0.05 for Welch Two Sample t-test.

**Figure S4 (related to Fig. 4)**

A. hCEC clones with different aneuploidies were analyzed by bulk WGS and by scRNAseq. Arm-level copy number events were inferred for WGS and for scRNAseq (see also **Methods**) and the deriving copy number profiles are shown for both methods. See also Fig. S4B.
B. hCEC clones containing trisomy of chr 7 or more complex karyotype were analyzed by FISH or by scRNA seq and the percentage of aneuploid cells was quantified using both methods. See text and Fig. S4A for additional details.
C. A heatmap depicting gene copy numbers inferred from scRNAseq analysis following KaryoCreate control experiments. hCEC cells were transduced either with empty vector or with KaryoCreate vector together with a negative control sgRNA (sgRNA NC) and scRNAseq was performed as in Fig. 4B to estimate % of gains and losses across chromosomes.
D. A heatmap depicting gene copy numbers inferred from scRNAseq analysis following KaryoCreate. KaryoCreate for different individual chromosomes (or combination of chromosomes) was performed on RPE-1 cells. scRNAseq was used to estimate the presence of chromosome or arm-level gains or losses using a modified version of CopyKat. The median expression of genes across each chromosome arm is used to estimate the DNA copy number. The % of gains and losses for each arm (reported below each heatmap) is estimated by comparing the DNA copy number distribution of each experimental sample (chromosome-specific sgRNA) to the negative control (sgRNA NC; see also **Methods**). In the heatmaps the rows represent individual cells, the columns represent different chromosomes, and the color represents the copy number change (gain in red and loss in blue).
E. Average proportion (%) of whole chromosome gains/losses and arm-level gains and losses. The percentage of the indicated events were calculated as the average among the aneuploid cells generated using KaryoCreate for chromosomes 6, 7, 8, 9, 11, 12 and X.
F. A heatmap depicting gene copy numbers inferred from scRNAseq analysis following KaryoCreate. KaryoCreate for different individual chromosomes (or combination of chromosomes) was performed on hCEC cells using two sgRNAs targeting chromosome 7 (sgChr7-1) and 18 (sgChr18-4). scRNAseq was used to estimate the presence of chromosome or arm-level gains or losses using a modified version of CopyKat. The median expression of genes across each chromosome arm is used to estimate the DNA copy number. The % of gains and losses for each arm (reported below each heatmap) is estimated by comparing the DNA copy number distribution of each experimental sample (chromosome-specific sgRNA) to the negative control (sgRNA NC; see also **Methods**). In the heatmaps the rows represent individual cells, the columns represent different chromosomes, and the color represents the copy number change (gain in red and loss in blue).

**Figure S5 (related to Fig. 5)**

A. Schematics of experimental plan involving KaryoCreate across different chromosomes to derive single cell clones with chromosome specific gains or losses.
B. KaryoCreate using a NC sgRNA, or sgRNA specific for chr7 (as indicated) was performed on hCEC hTERT p53-/- diploid cells and single cell-derived clones were analyzed by shallow WGS.
C. Survival analysis (Kaplan-Meier curve) for colorectal cancer patients ((TCGA – COADREAD) displaying or not 18q loss after excluding patients with SMAD4 point mutation.
D. Proliferation rate for clones 13 and 14 (18q loss) (as in Fig. 5E) after the overexpression of the indicated genes. Mean and standard deviation are shown for triplicates; p-values are from Wilcoxon test (*=p<0.05).

## REFERENCES

Altemose, N., Logsdon, G.A., Bzikadze, A.V., Sidhwani, P., Langley, S.A., Caldas, G.V., Hoyt, S.J., Uralsky, L., Ryabov, F.D., Shew, C.J., et al. (2022). Complete genomic and epigenetic maps of human centromeres. Science 376, eabl4178. https://doi.org/10.1126/science.abl4178.

Banaszynski, L.A., Chen, L.-C., Maynard-Smith, L.A., Ooi, A.G.L., and Wandless, T.J. (2006). A rapid, reversible, and tunable method to regulate protein function in living cells using synthetic small molecules. Cell 126, 995–1004. https://doi.org/10.1016/j.cell.2006.07.025.

Barra, V., and Fachinetti, D. (2018). The dark side of centromeres: types, causes and consequences of structural abnormalities implicating centromeric DNA. Nat. Commun. 9, 4340. https://doi.org/10.1038/s41467-018-06545-y.

Beroukhim, R., Mermel, C.H., Porter, D., Wei, G., Raychaudhuri, S., Donovan, J., Barretina, J., Boehm, J.S., Dobson, J., Urashima, M., et al. (2010). The landscape of somatic copy-number alteration across human cancers. Nature 463, 899–905. https://doi.org/10.1038/nature08822.

Bury, L., Moodie, B., Ly, J., McKay, L.S., Miga, K.H., and Cheeseman, I.M. (2020). Alpha-satellite RNA transcripts are repressed by centromere-nucleolus associations. ELife 9, e59770. https://doi.org/10.7554/eLife.59770.

Cheeseman, I.M. (2014). The kinetochore. Cold Spring Harb. Perspect. Biol. 6, a015826. https://doi.org/10.1101/cshperspect.a015826.

Cimini, D., Howell, B., Maddox, P., Khodjakov, A., Degrassi, F., and Salmon, E.D. (2001). Merotelic kinetochore orientation is a major mechanism of aneuploidy in mitotic mammalian tissue cells. J. Cell Biol. 153, 517–527. https://doi.org/10.1083/jcb.153.3.517.

Davoli, T., Xu, A.W., Mengwasser, K.E., Sack, L.M., Yoon, J.C., Park, P.J., and Elledge, S.J. (2013). Cumulative haploinsufficiency and triplosensitivity drive aneuploidy patterns and shape the cancer genome. Cell 155, 948–962. https://doi.org/10.1016/j.cell.2013.10.011.

DeLuca, J.G., Gall, W.E., Ciferri, C., Cimini, D., Musacchio, A., and Salmon, E.D. (2006). Kinetochore microtubule dynamics and attachment stability are regulated by Hec1. Cell 127, 969–982. https://doi.org/10.1016/j.cell.2006.09.047.

Dobin, A., Davis, C.A., Schlesinger, F., Drenkow, J., Zaleski, C., Jha, S., Batut, P., Chaisson, M., and Gingeras, T.R. (2013). STAR: ultrafast universal RNA-seq aligner. Bioinforma. Oxf. Engl. 29, 15–21. https://doi.org/10.1093/bioinformatics/bts635.

Doench, J.G., Hartenian, E., Graham, D.B., Tothova, Z., Hegde, M., Smith, I., Sullender, M., Ebert, B.L., Xavier, R.J., and Root, D.E. (2014). Rational design of highly active sgRNAs for CRISPR-Cas9-mediated gene inactivation. Nat. Biotechnol. 32, 1262–1267. https://doi.org/10.1038/nbt.3026.

Doench, J.G., Fusi, N., Sullender, M., Hegde, M., Vaimberg, E.W., Donovan, K.F., Smith, I., Tothova, Z., Wilen, C., Orchard, R., et al. (2016). Optimized sgRNA design to maximize activity and minimize off-target effects of CRISPR-Cas9. Nat. Biotechnol. 34, 184–191. https://doi.org/10.1038/nbt.3437.

Dolgalev, Igor (2022). Seq-N-Slide (Zenodo).

Drost, J., van Jaarsveld, R.H., Ponsioen, B., Zimberlin, C., van Boxtel, R., Buijs, A., Sachs, N., Overmeer, R.M., Offerhaus, G.J., Begthel, H., et al. (2015). Sequential cancer mutations in cultured human intestinal stem cells. Nature 521, 43–47. https://doi.org/10.1038/nature14415.

Dumont, M., Gamba, R., Gestraud, P., Klaasen, S., Worrall, J.T., De Vries, S.G., Boudreau, V., Salinas-Luypaert, C., Maddox, P.S., Lens, S.M., et al. (2020). Human chromosome-specific aneuploidy is influenced by DNA -dependent centromeric features. EMBO J. 39. https://doi.org/10.15252/embj.2019102924.

Eppert, K., Scherer, S.W., Ozcelik, H., Pirone, R., Hoodless, P., Kim, H., Tsui, L.C., Bapat, B., Gallinger, S., Andrulis, I.L., et al. (1996). MADR2 maps to 18q21 and encodes a TGFbeta-regulated MAD-related protein that is functionally mutated in colorectal carcinoma. Cell 86, 543–552. https://doi.org/10.1016/s0092-8674(00)80128-2.

Fournier, R.E. (1981). A general high-efficiency procedure for production of microcell hybrids. Proc. Natl. Acad. Sci. U. S. A. 78, 6349–6353. https://doi.org/10.1073/pnas.78.10.6349.

Gao, R., Bai, S., Henderson, Y.C., Lin, Y., Schalck, A., Yan, Y., Kumar, T., Hu, M., Sei, E., Davis, A., et al. (2021). Delineating copy number and clonal substructure in human tumors from single-cell transcriptomes. Nat. Biotechnol. 39, 599–608. https://doi.org/10.1038/s41587-020-00795-2.

Gordon, D.J., Resio, B., and Pellman, D. (2012). Causes and consequences of aneuploidy in cancer. Nat. Rev. Genet. 13, 189–203. https://doi.org/10.1038/nrg3123.

Gregan, J., Polakova, S., Zhang, L., Tolić-Nørrelykke, I.M., and Cimini, D. (2011). Merotelic kinetochore attachment: causes and effects. Trends Cell Biol. 21, 374–381. https://doi.org/10.1016/j.tcb.2011.01.003.

Groth, K.A., Skakkebæk, A., Høst, C., Gravholt, C.H., and Bojesen, A. (2013). Clinical review: Klinefelter syndrome--a clinical update. J. Clin. Endocrinol. Metab. 98, 20–30. https://doi.org/2016092613072900074.

Hayden, K.E. (2012). Human centromere genomics: now it’s personal. Chromosome Res. Int. J. Mol. Supramol. Evol. Asp. Chromosome Biol. 20, 621–633. https://doi.org/10.1007/s10577-012-9295-y.

Kirby, R.S. (2017). The prevalence of selected major birth defects in the United States. Semin. Perinatol. 41, 338–344. https://doi.org/10.1053/j.semperi.2017.07.004.

Knouse, K.A., Wu, J., Whittaker, C.A., and Amon, A. (2014). Single cell sequencing reveals low levels of aneuploidy across mammalian tissues. Proc. Natl. Acad. Sci. U. S. A. 111, 13409–13414. https://doi.org/10.1073/pnas.1415287111.

Knouse, K.A., Davoli, T., Elledge, S.J., and Amon, A. (2017). Aneuploidy in Cancer: Seq-ing Answers to Old Questions. Annu. Rev. Cancer Biol. 1, 335–354. https://doi.org/10.1146/annurev-cancerbio-042616-072231.

Korenbaum, E., and Rivero, F. (2002). Calponin homology domains at a glance. J. Cell Sci. 115, 3543–3545. https://doi.org/10.1242/jcs.00003.

Kuilman, T., Velds, A., Kemper, K., Ranzani, M., Bombardelli, L., Hoogstraat, M., Nevedomskaya, E., Xu, G., de Ruiter, J., Lolkema, M.P., et al. (2015). CopywriteR: DNA copy number detection from off-target sequence data. Genome Biol. 16, 49. https://doi.org/10.1186/s13059-015-0617-1.

Li, H., and Durbin, R. (2009). Fast and accurate short read alignment with Burrows-Wheeler transform. Bioinforma. Oxf. Engl. 25, 1754–1760. https://doi.org/10.1093/bioinformatics/btp324.

Liao, Y., Smyth, G.K., and Shi, W. (2014). featureCounts: an efficient general purpose program for assigning sequence reads to genomic features. Bioinforma. Oxf. Engl. 30, 923–930. https://doi.org/10.1093/bioinformatics/btt656.

Liu, D., Vleugel, M., Backer, C.B., Hori, T., Fukagawa, T., Cheeseman, I.M., and Lampson, M.A. (2010). Regulated targeting of protein phosphatase 1 to the outer kinetochore by KNL1 opposes Aurora B kinase. J. Cell Biol. 188, 809–820. https://doi.org/10.1083/jcb.201001006.

Liu, J., Lichtenberg, T., Hoadley, K.A., Poisson, L.M., Lazar, A.J., Cherniack, A.D., Kovatich, A.J., Benz, C.C., Levine, D.A., Lee, A.V., et al. (2018). An Integrated TCGA Pan-Cancer Clinical Data Resource to Drive High-Quality Survival Outcome Analytics. Cell 173, 400–416.e11. https://doi.org/10.1016/j.cell.2018.02.052.

Love, M.I., Huber, W., and Anders, S. (2014). Moderated estimation of fold change and dispersion for RNA-seq data with DESeq2. Genome Biol. 15, 550. https://doi.org/10.1186/s13059-014-0550-8.

Ly, P., Eskiocak, U., Kim, S.B., Roig, A.I., Hight, S.K., Lulla, D.R., Zou, Y.S., Batten, K., Wright, W.E., and Shay, J.W. (2011). Characterization of aneuploid populations with trisomy 7 and 20 derived from diploid human colonic epithelial cells. Neoplasia N. Y. N 13, 348–357. https://doi.org/10.1593/neo.101580.

Maciejowski, J., Li, Y., Bosco, N., Campbell, P.J., and de Lange, T. (2015). Chromothripsis and Kataegis Induced by Telomere Crisis. Cell 163, 1641–1654. https://doi.org/10.1016/j.cell.2015.11.054.

Maloney, K.A., Sullivan, L.L., Matheny, J.E., Strome, E.D., Merrett, S.L., Ferris, A., and Sullivan, B.A. (2012). Functional epialleles at an endogenous human centromere. Proc. Natl. Acad. Sci. U. S. A. 109, 13704–13709. https://doi.org/10.1073/pnas.1203126109.

Massagué, J., Blain, S.W., and Lo, R.S. (2000). TGFbeta signaling in growth control, cancer, and heritable disorders. Cell 103, 295–309. https://doi.org/10.1016/s0092-8674(00)00121-5.

McFadden, D.G., Papagiannakopoulos, T., Taylor-Weiner, A., Stewart, C., Carter, S.L., Cibulskis, K., Bhutkar, A., McKenna, A., Dooley, A., Vernon, A., et al. (2014). Genetic and clonal dissection of murine small cell lung carcinoma progression by genome sequencing. Cell 156, 1298–1311. https://doi.org/10.1016/j.cell.2014.02.031.

Meerbrey, K.L., Hu, G., Kessler, J.D., Roarty, K., Li, M.Z., Fang, J.E., Herschkowitz, J.I., Burrows, A.E., Ciccia, A., Sun, T., et al. (2011). The pINDUCER lentiviral toolkit for inducible RNA interference in vitro and in vivo. Proc. Natl. Acad. Sci. U. S. A. 108, 3665–3670. https://doi.org/10.1073/pnas.1019736108.

Musacchio, A. (2015). The Molecular Biology of Spindle Assembly Checkpoint Signaling Dynamics. Curr. Biol. CB 25, R1002–1018. https://doi.org/10.1016/j.cub.2015.08.051.

Musacchio, A., and Desai, A. (2017). A Molecular View of Kinetochore Assembly and Function. Biology 6, E5. https://doi.org/10.3390/biology6010005.

Nurk, S., Koren, S., Rhie, A., Rautiainen, M., Bzikadze, A.V., Mikheenko, A., Vollger, M.R., Altemose, N., Uralsky, L., Gershman, A., et al. (2022). The complete sequence of a human genome. Science 376, 44–53. https://doi.org/10.1126/science.abj6987.

Patel, A.P., Tirosh, I., Trombetta, J.J., Shalek, A.K., Gillespie, S.M., Wakimoto, H., Cahill, D.P., Nahed, B.V., Curry, W.T., Martuza, R.L., et al. (2014). Single-cell RNA-seq highlights intratumoral heterogeneity in primary glioblastoma. Science 344, 1396–1401. https://doi.org/10.1126/science.1254257.

Ran, F.A., Hsu, P.D., Wright, J., Agarwala, V., Scott, D.A., and Zhang, F. (2013). Genome engineering using the CRISPR-Cas9 system. Nat. Protoc. 8, 2281–2308. https://doi.org/10.1038/nprot.2013.143.

Rayner, E., Durin, M.-A., Thomas, R., Moralli, D., O’Cathail, S.M., Tomlinson, I., Green, C.M., and Lewis, A. (2019). CRISPR-Cas9 Causes Chromosomal Instability and Rearrangements in Cancer Cell Lines, Detectable by Cytogenetic Methods. CRISPR J. 2, 406–416. https://doi.org/10.1089/crispr.2019.0006.

Sanjana, N.E., Shalem, O., and Zhang, F. (2014). Improved vectors and genome-wide libraries for CRISPR screening. Nat. Methods 11, 783–784. https://doi.org/10.1038/nmeth.3047.

Santaguida, S., and Amon, A. (2015). Short- and long-term effects of chromosome mis-segregation and aneuploidy. Nat. Rev. Mol. Cell Biol. 16, 473–485. https://doi.org/10.1038/nrm4025.

Schneider, V.A., Graves-Lindsay, T., Howe, K., Bouk, N., Chen, H.-C., Kitts, P.A., Murphy, T.D., Pruitt, K.D., Thibaud-Nissen, F., Albracht, D., et al. (2017). Evaluation of GRCh38 and de novo haploid genome assemblies demonstrates the enduring quality of the reference assembly. Genome Res. 27, 849–864. https://doi.org/10.1101/gr.213611.116.

Schueler, M.G., and Sullivan, B.A. (2006). Structural and functional dynamics of human centromeric chromatin. Annu. Rev. Genomics Hum. Genet. 7, 301–313. https://doi.org/10.1146/annurev.genom.7.080505.115613.

Sheppard, O., Wiseman, F.K., Ruparelia, A., Tybulewicz, V.L.J., and Fisher, E.M.C. (2012). Mouse Models of Aneuploidy. Sci. World J. 2012, 1–6. https://doi.org/10.1100/2012/214078.

Stern, B.M., and Murray, A.W. (2001). Lack of tension at kinetochores activates the spindle checkpoint in budding yeast. Curr. Biol. CB 11, 1462–1467. https://doi.org/10.1016/s0960-9822(01)00451-1.

Stingele, S., Stoehr, G., Peplowska, K., Cox, J., Mann, M., and Storchova, Z. (2012). Global analysis of genome, transcriptome and proteome reveals the response to aneuploidy in human cells. Mol. Syst. Biol. 8, 608. https://doi.org/10.1038/msb.2012.40.

Subramanian, A., Tamayo, P., Mootha, V.K., Mukherjee, S., Ebert, B.L., Gillette, M.A., Paulovich, A., Pomeroy, S.L., Golub, T.R., Lander, E.S., et al. (2005). Gene set enrichment analysis: a knowledge-based approach for interpreting genome-wide expression profiles. Proc. Natl. Acad. Sci. U. S. A. 102, 15545–15550. https://doi.org/10.1073/pnas.0506580102.

Sullivan, L.L., and Sullivan, B.A. (2020). Genomic and functional variation of human centromeres. Exp. Cell Res. 389, 111896. https://doi.org/10.1016/j.yexcr.2020.111896.

Taylor, A.M., Shih, J., Ha, G., Gao, G.F., Zhang, X., Berger, A.C., Schumacher, S.E., Wang, C., Hu, H., Liu, J., et al. (2018). Genomic and Functional Approaches to Understanding Cancer Aneuploidy. Cancer Cell 33, 676–689.e3. https://doi.org/10.1016/j.ccell.2018.03.007.

Thiagalingam, S., Lengauer, C., Leach, F.S., Schutte, M., Hahn, S.A., Overhauser, J., Willson, J.K., Markowitz, S., Hamilton, S.R., Kern, S.E., et al. (1996). Evaluation of candidate tumour suppressor genes on chromosome 18 in colorectal cancers. Nat. Genet. 13, 343–346. https://doi.org/10.1038/ng0796-343.

Tirosh, I., Izar, B., Prakadan, S.M., Wadsworth, M.H., Treacy, D., Trombetta, J.J., Rotem, A., Rodman, C., Lian, C., Murphy, G., et al. (2016). Dissecting the multicellular ecosystem of metastatic melanoma by single-cell RNA-seq. Science 352, 189–196. https://doi.org/10.1126/science.aad0501.

Tovini, L., Johnson, S.C., Andersen, A.M., Spierings, D.C.J., Wardenaar, R., Foijer, F., and McClelland, S.E. (2022). Inducing Specific Chromosome Mis-Segregation in Human Cells. 2022.04.19.486691. https://doi.org/10.1101/2022.04.19.486691.

Trakala, M., Aggarwal, M., Sniffen, C., Zasadil, L., Carroll, A., Ma, D., Su, X.A., Wangsa, D., Meyer, A., Sieben, C.J., et al. (2021). Clonal selection of stable aneuploidies in progenitor cells drives high-prevalence tumorigenesis. Genes Dev. 35, 1079–1092. https://doi.org/10.1101/gad.348341.121.

Truong, M.A., Cané-Gasull, P., Vries, S.G. de, Nijenhuis, W., Wardenaar, R., Kapitein, L.C., Foijer, F., and Lens, S.M.A. (2022). A motor-based approach to induce chromosome-specific mis-segregations in human cells. 2022.04.19.488790. https://doi.org/10.1101/2022.04.19.488790.

Uralsky, L.I., Shepelev, V.A., Alexandrov, A.A., Yurov, Y.B., Rogaev, E.I., and Alexandrov, I.A. (2019). Classification and monomer-by-monomer annotation dataset of suprachromosomal family 1 alpha satellite higher-order repeats in hg38 human genome assembly. Data Brief 24, 103708. https://doi.org/10.1016/j.dib.2019.103708.

van der Walt, S., Schönberger, J.L., Nunez-Iglesias, J., Boulogne, F., Warner, J.D., Yager, N., Gouillart, E., Yu, T., and scikit-image contributors (2014). scikit-image: image processing in Python. PeerJ 2, e453. https://doi.org/10.7717/peerj.453.

Wang, T., Wei, J.J., Sabatini, D.M., and Lander, E.S. (2014). Genetic screens in human cells using the CRISPR-Cas9 system. Science 343, 80–84. https://doi.org/10.1126/science.1246981.

Watkins, T.B.K., Lim, E.L., Petkovic, M., Elizalde, S., Birkbak, N.J., Wilson, G.A., Moore, D.A., Grönroos, E., Rowan, A., Dewhurst, S.M., et al. (2020). Pervasive chromosomal instability and karyotype order in tumour evolution. Nature 587, 126–132. https://doi.org/10.1038/s41586-020-2698-6.

van de Wetering, M., Francies, H.E., Francis, J.M., Bounova, G., Iorio, F., Pronk, A., van Houdt, W., van Gorp, J., Taylor-Weiner, A., Kester, L., et al. (2015). Prospective derivation of a living organoid biobank of colorectal cancer patients. Cell 161, 933–945. https://doi.org/10.1016/j.cell.2015.03.053.

Whinn, K.S., Kaur, G., Lewis, J.S., Schauer, G.D., Mueller, S.H., Jergic, S., Maynard, H., Gan, Z.Y., Naganbabu, M., Bruchez, M.P., et al. (2019). Nuclease dead Cas9 is a programmable roadblock for DNA replication. Sci. Rep. 9, 13292. https://doi.org/10.1038/s41598-019-49837-z.

Willard, H.F. (1991). Evolution of alpha satellite. Curr. Opin. Genet. Dev. 1, 509–514. https://doi.org/10.1016/s0959-437x(05)80200-x.

William, W.N., Zhao, X., Bianchi, J.J., Lin, H.Y., Cheng, P., Lee, J.J., Carter, H., Alexandrov, L.B., Abraham, J.P., Spetzler, D.B., et al. (2021). Immune evasion in HPV-head and neck precancer-cancer transition is driven by an aneuploid switch involving chromosome 9p loss. Proc. Natl. Acad. Sci. U. S. A. 118, e2022655118. https://doi.org/10.1073/pnas.2022655118.

Xue, W., Kitzing, T., Roessler, S., Zuber, J., Krasnitz, A., Schultz, N., Revill, K., Weissmueller, S., Rappaport, A.R., Simon, J., et al. (2012). A cluster of cooperating tumor-suppressor gene candidates in chromosomal deletions. Proc. Natl. Acad. Sci. U. S. A. 109, 8212–8217. https://doi.org/10.1073/pnas.1206062109.

Zuo, E., Huo, X., Yao, X., Hu, X., Sun, Y., Yin, J., He, B., Wang, X., Shi, L., Ping, J., et al. (2017). CRISPR/Cas9-mediated targeted chromosome elimination. Genome Biol. 18, 224. https://doi.org/10.1186/s13059-017-1354-4.

