## Supplementary figures and images for "KaryoCreate: a new CRISPR-based technology to generate chromosome-specific aneuploidy by targeting human centromeres"

### Figure S1

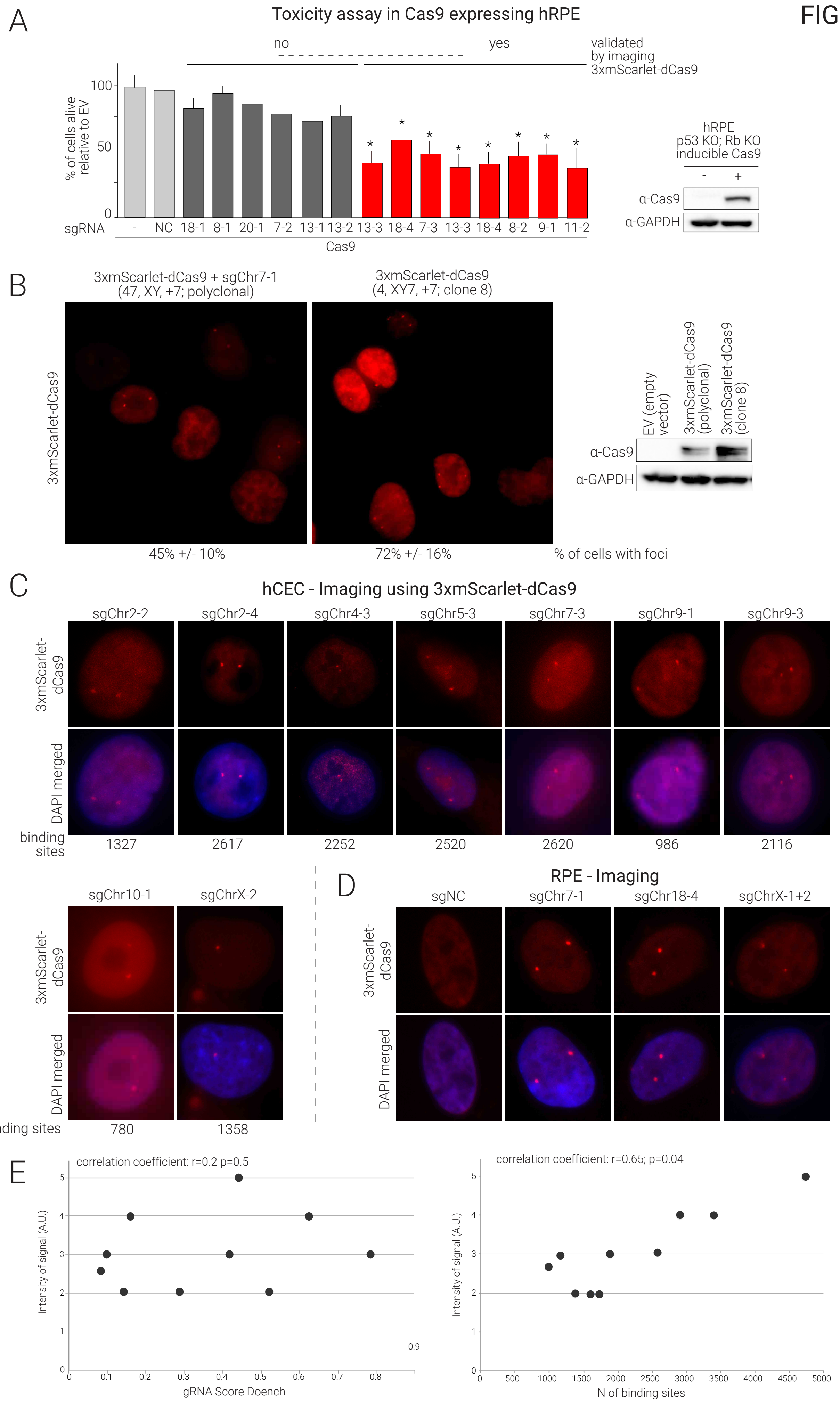

### Figure S2

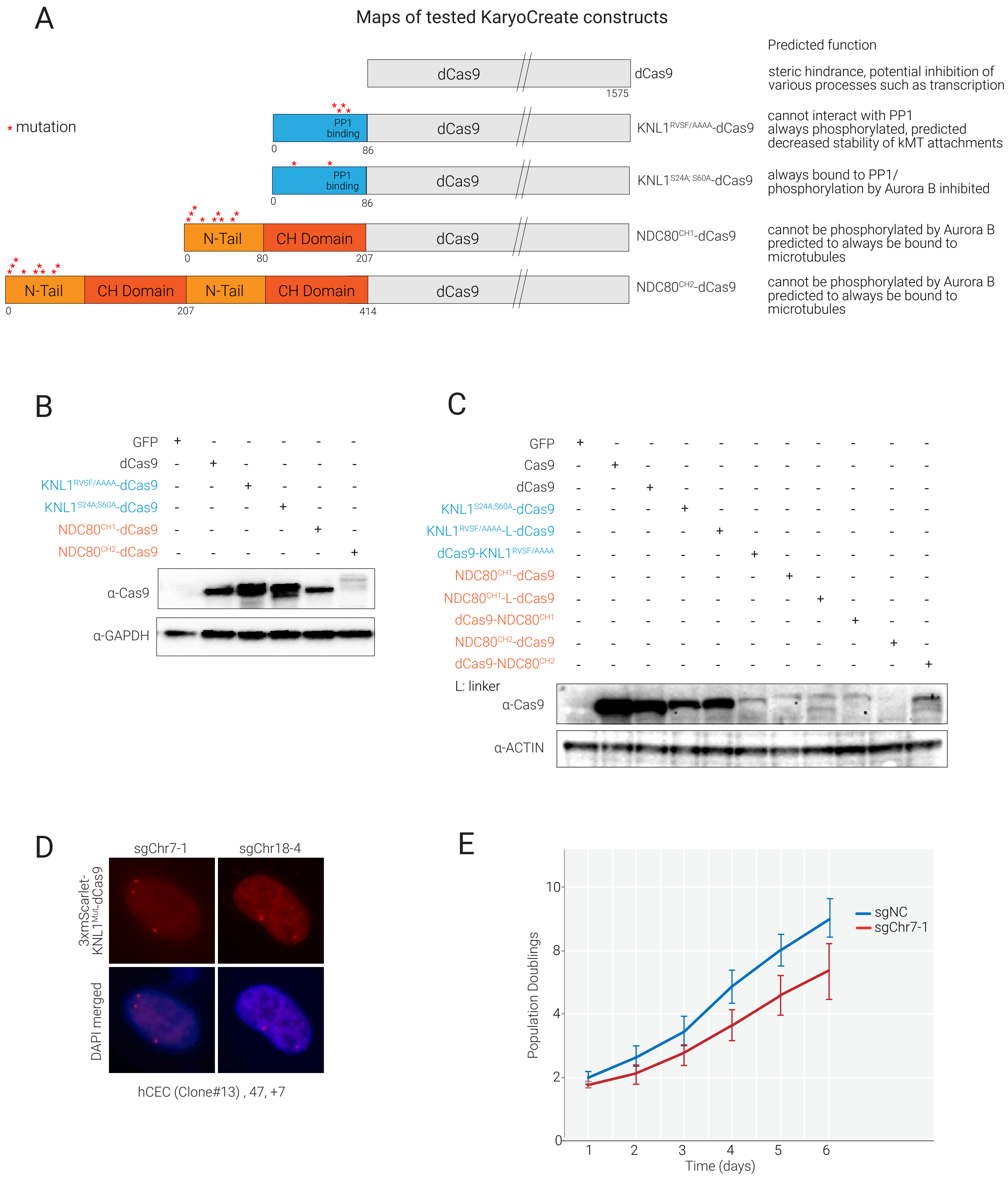

### Figure S3

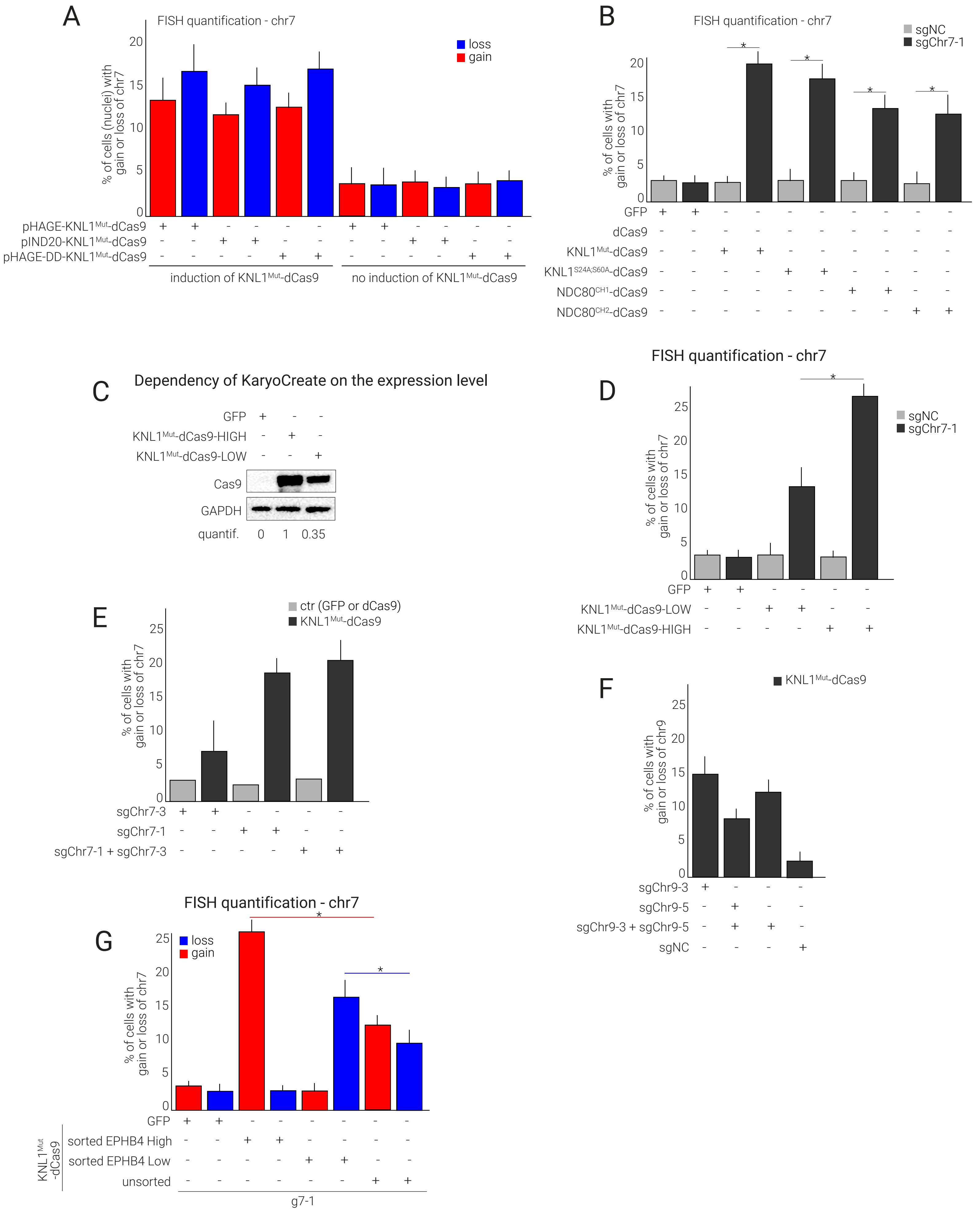

### Figure S4

#### FIGURE S4

##### Validation of scRNAseq to infer aneuploidy on aneuploid clones

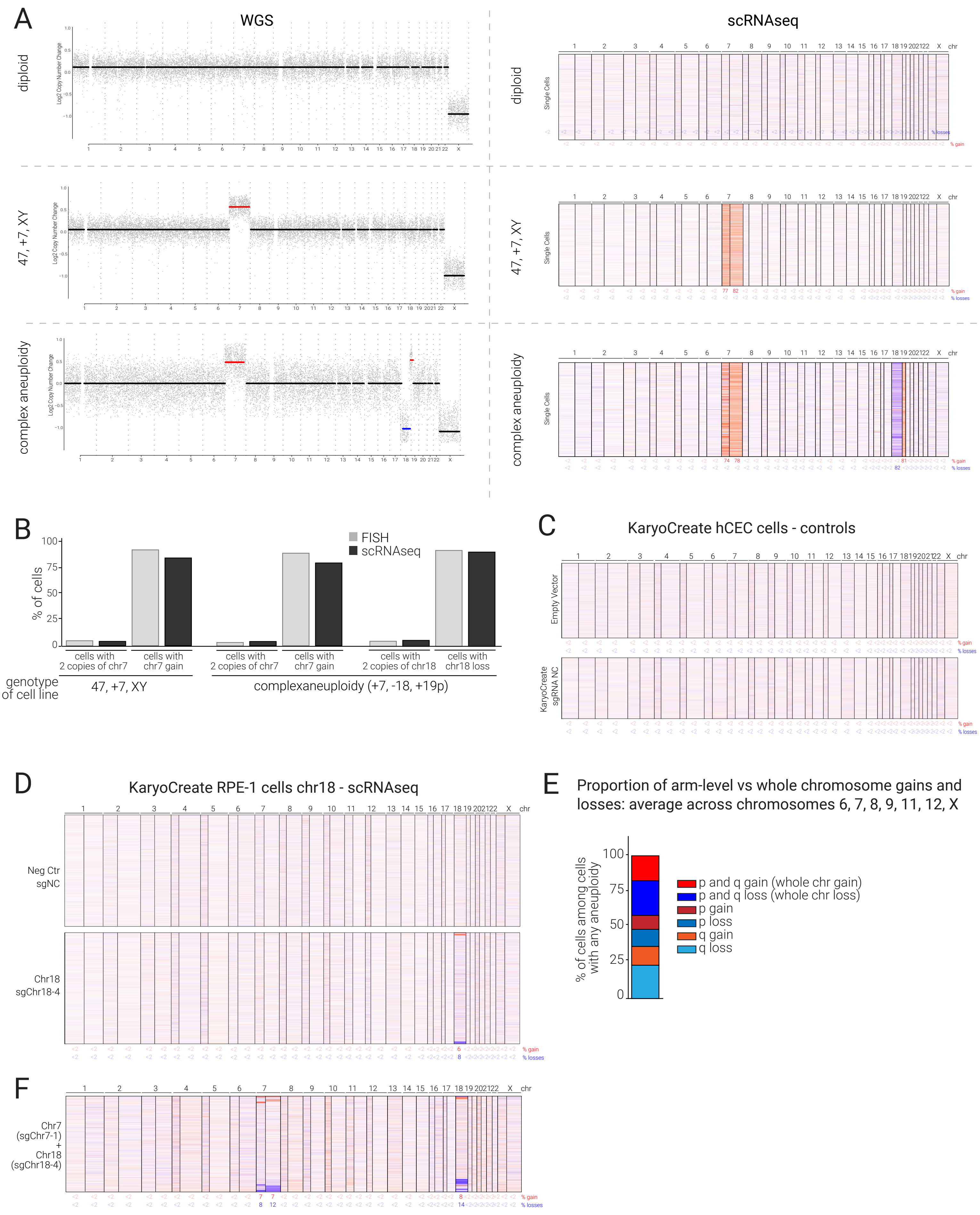

### Figure S5

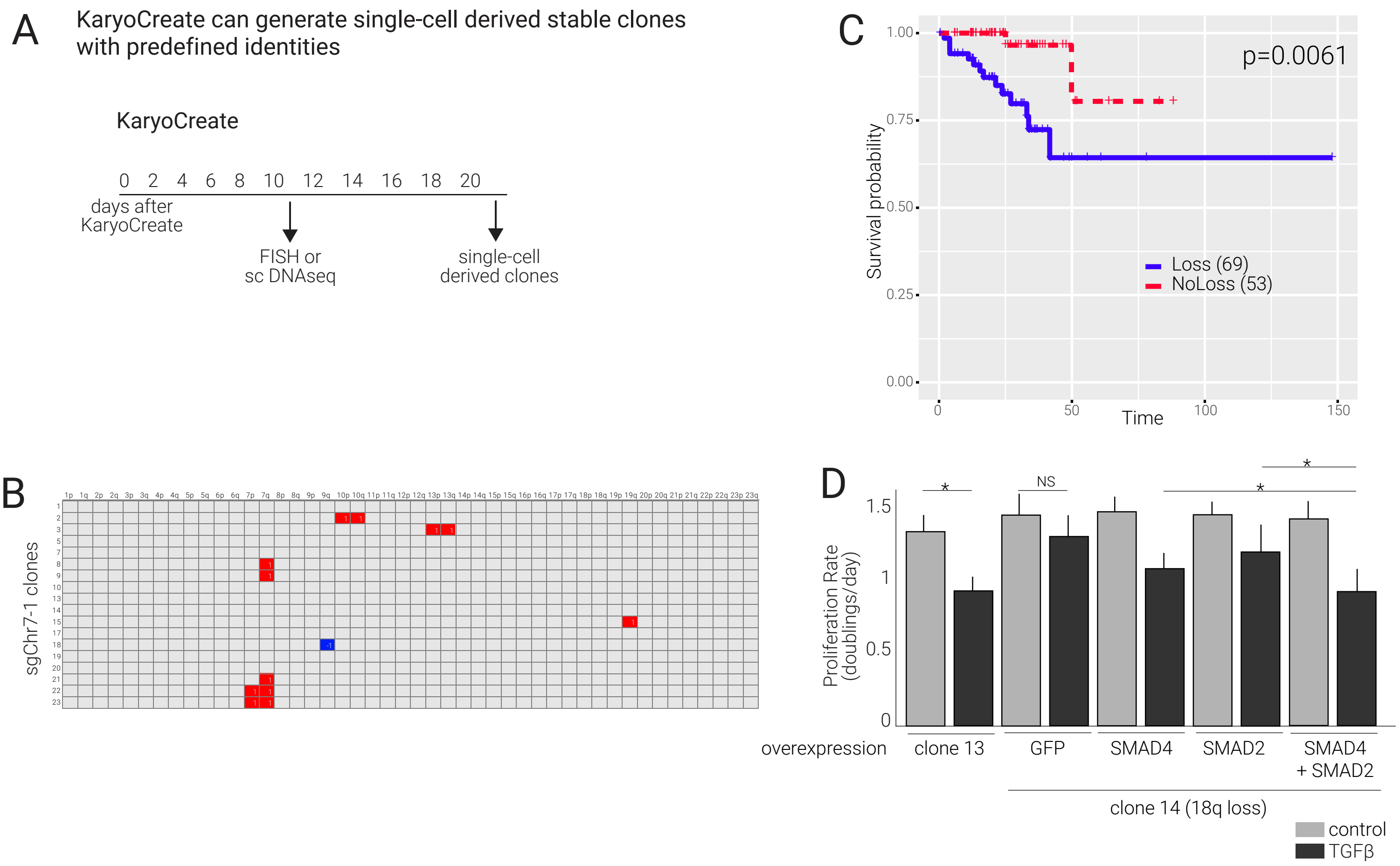
